# Dietary protein shapes the profile and repertoire of intestinal CD4^+^ T cells

**DOI:** 10.1101/2023.04.11.536475

**Authors:** Ainsley Lockhart, Aubrey Reed, Tiago Rezende de Castro, Calvin Herman, Maria Cecilia Campos Canesso, Daniel Mucida

## Abstract

The intestinal immune system must tolerate food antigens to avoid allergy, a process requiring CD4^+^ T cells. Combining antigenically defined diets with gnotobiotic models, we show that food and microbiota distinctly influence the profile and T cell receptor repertoire of intestinal CD4^+^ T cells. Independent of the microbiota, dietary proteins contributed to accumulation and clonal selection of antigen-experienced CD4^+^ T cells at the intestinal epithelium, imprinting a tissue specialized transcriptional program including cytotoxic genes on both conventional and regulatory CD4^+^ T cells (Tregs). This steady state CD4^+^ T cell response to food was disrupted by inflammatory challenge, and protection against food allergy in this context was associated with Treg clonal expansion and decreased pro-inflammatory gene expression. Finally, we identified both steady state epithelium-adapted CD4^+^ T cells and tolerance-induced Tregs that recognize dietary antigens, suggesting that both cell types may be critical for preventing inappropriate immune responses to food.

## Introduction

Large quantities of food-derived antigens are absorbed through the intestine each day which must be tolerated by the immune system to avoid food allergies. Oral tolerance, a key mechanism whereby oral administration of antigen results in both local and systemic tolerance to that antigen, requires CD4^+^ T cells including specifically regulatory T cells (Tregs) (Garside et al., 1995; Hadis et al., 2011; Josefowicz et al., 2012; Mucida et al., 2005; Pabst and Mowat, 2012). However, T cell responses to dietary antigen have primarily been characterized using monoclonal T cell receptor (TCR) transgenic systems which do not represent a physiological immune response. Polyclonal CD4^+^ T cell responses to food, including TCR-specific selection and functional differentiation, remain largely uncharacterized and are critical for understanding mechanisms of tolerance and allergy.

CD4^+^ T cells occupy two major adjacent tissue compartments in the intestine, the lamina propria (LP) and epithelium (IE), which are segregated by a basement membrane and are immunologically distinct. Tregs, which are critical mediators of intestinal inflammation, are enriched in the LP but relatively rare in the highly selective IE (Sujino et al., 2016). Upon migration to the IE, both conventional CD4^+^ T cells and Tregs can undergo stepwise acquisition of a specialized transcriptional program, upregulating genes associated with tissue residency (CD103, CD69), cytotoxicity (granzymes), natural killer function (NKG7), and CD8^+^ T cell lineage (Runx3) (London et al., 2021; Mucida et al., 2013; Reis et al., 2013; Sujino et al., 2016). At the terminal point of this differentiation process, IE-adapted CD4^+^ T cells upregulate the CD8αα homodimer, which is proposed to dampen TCR signaling resulting in a high activation threshold. (Cheroutre and Lambolez, 2008). Our recent findings suggest that IE-adapted CD4^+^ T cells play a complementary anti-inflammatory role to Tregs, providing an important regulatory mechanism in the gut epithelium where Tregs are rare (Bilate et al., 2016; Bilate et al., 2020; Bousbaine et al., 2022; Sujino et al., 2016). Disruption of Treg generation (Bouziat et al., 2017; Josefowicz et al., 2012; Torgerson et al., 2007) or epithelial T cell programming (Reis et al., 2013; Sujino et al., 2016) leads to intestinal inflammation. Additionally, IE-adapted CD4^+^ T cells can contribute to immune regulation towards dietary antigen (Sujino et al., 2016). However, IE-adapted CD4^+^ T cells also have pro-inflammatory potential and can play a pathological role in a dysregulated response to dietary antigen as is seen in Celiac disease (Abadie et al., 2012; Abadie et al., 2020; Costes et al., 2019; Fina et al., 2008).

Here, we characterize polyclonal intestinal CD4^+^ T cell responses to dietary protein and demonstrate that their prevalent steady state fate in the intestine is acquisition of an epithelium residency-associated transcriptional profile including expression of cytotoxicity-associated genes. We further demonstrate how intestinal T cell responses to food are altered in active tolerance or allergy to favor tissue influx of Tregs or pro-inflammatory T helper cells, respectively. These findings suggest that epithelium-adapted CD4^+^ T cells in addition to Tregs contribute to homeostatic immune responses to food.

## Results

### Dietary signals promote accumulation of mature CD4^+^ T cells in the gut epithelium

To characterize intestinal T cell responses to food protein, we developed a protein antigen-free solid diet (AA) containing free amino acids. AA diet lacks polypeptides but does contain other dietary macromolecules including simple and complex carbohydrates (sucrose, corn starch), lipids (corn oil), and fiber (cellulose) **(Table S1)**. Standard chow diet by contrast is highly complex, comprised of whole food ingredients (e.g., wheat, corn, soybeans) which contain hundreds of distinct proteins and diverse dietary metabolites. Chow diet can therefore impact intestinal T cells through dietary antigen-specific TCR stimulation, immune co-stimulation by dietary metabolites (some in a microbiota-dependent manner), and secondary stimulation via diet-induced changes to the microbiota.

Specific pathogen-free (SPF) C57BL/6 mice weaned onto AA diet were similar to standard chow diet mice in weight gain, intestinal inflammation measured by fecal lipocalin-2, and serum nutritional biomarkers **(Figure S1A-B).** Although the length of the small intestine was reduced in 8-week-old AA diet mice **(Figure S1C),** histological examination revealed no evidence of tissue damage or inflammation and minimal differences in tissue architecture **(Figures S1D-E).** Finally, we found no significant differences in intestinal myeloid cell frequencies in SPF AA vs chow diet mice **(Figure S1F)**. Altogether, AA diet mice appear heathy and display no evidence of intestinal damage or inflammation.

We assessed the two major intestinal T cell compartments, the IE and LP, by flow cytometry, and found increased TCRαβ^+^ CD4^+^ T cells in the small intestine IE of 8-week-old mice weaned onto chow died compared to AA diet **(Figure 1A)**. Total TCRαβ^+^ T cells were also increased in the small intestine LP of chow diet mice; however, chow diet did not promote increased frequency of CD4^+^ T cells in this compartment **(Figure S2A)**. Antigen-experienced CD44^+^ CD62L^-^ cells, including tissue adapted pre-CD8αα^+^ (CD103^+^ CD8αα^-^) and CD8αα^+^ subsets (London et al., 2021), accounted for the vast majority of IE CD4^+^ T cells induced by exposure to chow diet **(Figure 1B-C)**. Whereas the absolute number of IE Tregs was also slightly increased in chow diet, they were reduced by relative frequency out of CD4^+^, suggesting that exposure to dietary signals favors IE-adapted subsets over Tregs **(Figure 1B-C)**. Additionally, this may suggest that the epithelial conversion of Tregs into CD4^+^ CD8αα^+^ T cells may be impaired in AA-diet mice (Sujino et al., 2016).

**Figure 1.**
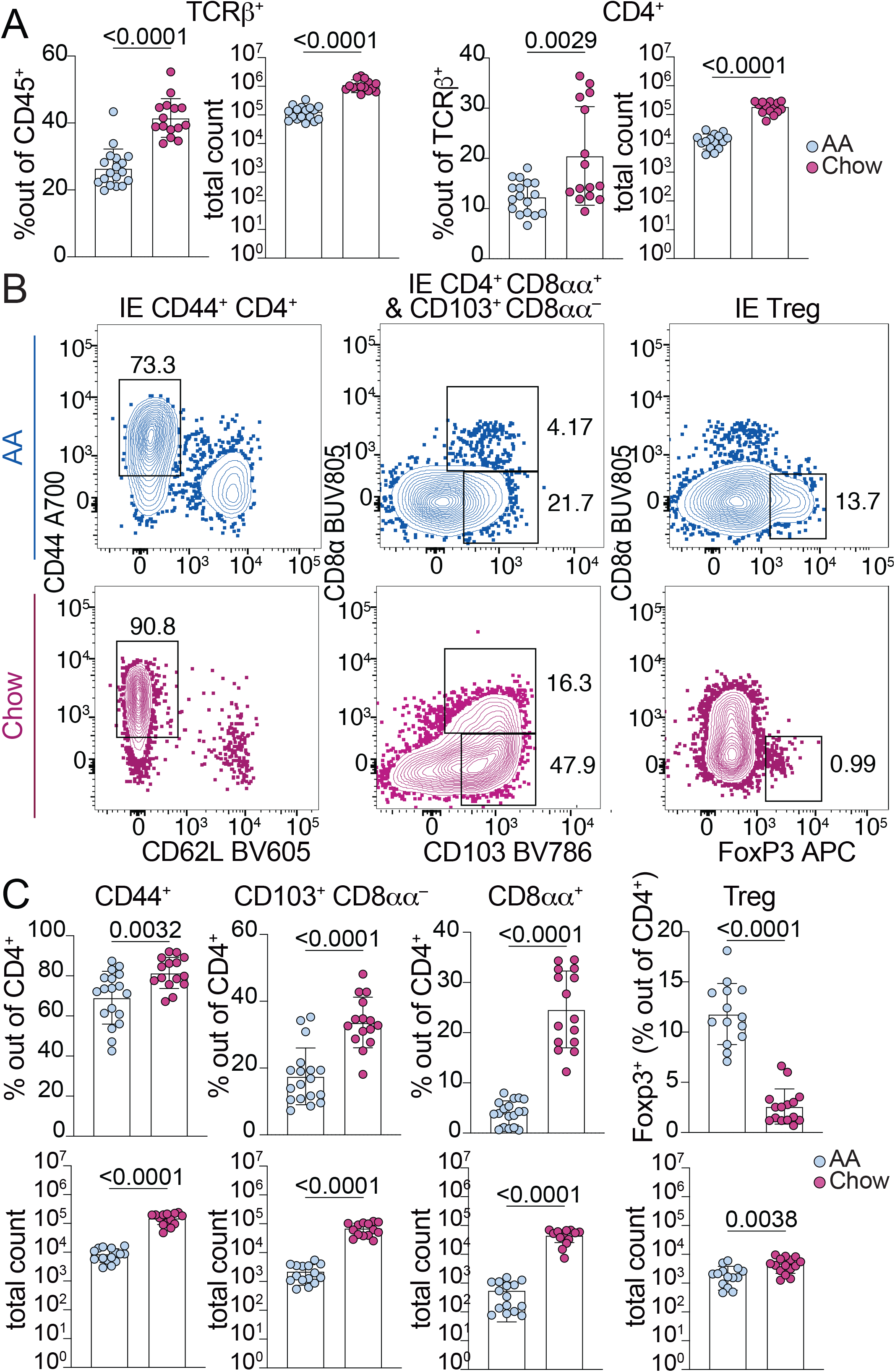
Dietary signals promote accumulation and adaptation of intestinal CD4+ T cells in the small intestine epithelium of specific pathogen-free mice. **(A-C)** Flow cytometry from the IE of SPF mice fed AA or standard chow diet measuring frequency or absolute count of the indicated cell subsets (A, C) or showing representative flow plots pre-gated on CD4+ T cells (B). Mean +/-SD from 3-5 independent experiments with 14-18 mice per condition. Unpaired t-tests showing p-values <0.05.

Among LP CD4^+^ T cells, chow diet mice had a slight increase in absolute number of CD44^+^ cells but no increase in Tregs compared to AA diet **(Figure S2B)**. Additionally, we did not find increased Rorγt^-^ peripherally-induced Tregs (pTregs), which reportedly respond to chow diet in a germ-free setting (Kim et al., 2016), in either the IE or LP **(Figure S2C)**. We expected to find the greatest impact of dietary protein in the small intestine, the primary site of food absorption. Consistent with this, we found no chow-associated increase in large intestine IE CD4^+^ T cells including CD44^+^ CD4^+^, pre-CD8αα^+^ (CD103^+^ CD8αα^-^), or Tregs, and only a small increase in CD4^+^ CD8αα^+^ T cells **(Figure S2D)**. Altogether, these data demonstrate that complex diet promotes maturation and epithelial adaptation of small intestine CD4^+^ T cells, whereas the large intestine and LP are relatively less affected, indicative of a distinct and localized T cell response to food.

To assess the impact of dietary antigen on T cell phenotypes in a highly controlled manner, we next supplemented AA diet with a low dose of OVA (0.1% in drinking water, ∼5 mg/day) intended to model the amount of a single food protein present in a protein-diverse diet. OVA supplied in drinking water at this dose for 2 days was sufficient to stimulate OVA-specific TCR transgenic CD4^+^ T cells *in vivo* in gut-draining lymph nodes **(Figure S2E)**. SPF C57BL/6 mice fed AA+OVA had slightly yet significantly elevated absolute counts of total IE CD4^+^ T cells, CD44^+^ CD4^+^ T cells, and CD4^+^ CD8αα^+^ T cells relative to AA diet mice though frequency was unchanged **(Figure S2F)**. We found no difference in pre-CD8αα^+^ (CD103^+^ CD8αα^-^), Tregs from the IE, or any tested LP T cell subsets **(Figure S2F-G)**. To address whether different or more diverse food proteins could further impact intestinal CD4^+^ T cells, we weaned SPF mice onto AA diets supplemented with either cow milk casein, or a mix of casein, gluten, and soy protein. Comparing frequencies of IE CD4^+^ CD8αα^+^ T cells and Tregs in the proximal small intestine, we found no difference between AA and casein, but increased CD4^+^ CD8αα^+^ T cells and decreased Tregs in casein-gluten-soy diet, suggesting that the Treg to CD4^+^ CD8αα^+^ differentiation pathway (Sujino et al., 2016) was induced in mice fed more diverse proteins **(Figure S2H)**. These results indicate that dietary protein can promote IE CD4^+^ T cell accumulation and maturation in SPF mice, and that increased diversity of dietary protein such as in casein-gluten-soy or standard chow diet may have an additive effect.

### Dietary signals promote microbiota-independent epithelial and cytotoxic programming of CD4^+^ T cells

Diet highly influences the gut microbiota, which can lead to indirect downstream effects on local immune cells (Sonnenburg and Backhed, 2016). Indeed, 16S rRNA sequencing of SPF AA versus chow diet mice revealed distinct gut microbiome composition, with chow diet promoting increased microbial diversity in both the small intestine and cecum **(Figure 2A-C, Figure S2I-J, Table S2)**. To better characterize microbiota-independent dietary impact on immune cells, we established our dietary models in germ-free (GF) mice and gnotobiotic mice colonized with Oligo-MM^12^, a stable, vertically transmissible consortium of 12 commensal strains representing members of the major bacterial phyla in the murine gut (Brugiroux et al., 2016). GF AA-diet mice lack exposure to foreign protein antigen and provide an ideal model to assess the impact of diet independent of the microbiota, while Oligo-MM^12^ provides a more physiological model with some co-stimulation from commensal bacteria while still limiting the diversity and complexity of intestinal antigen.

**Figure 2.**
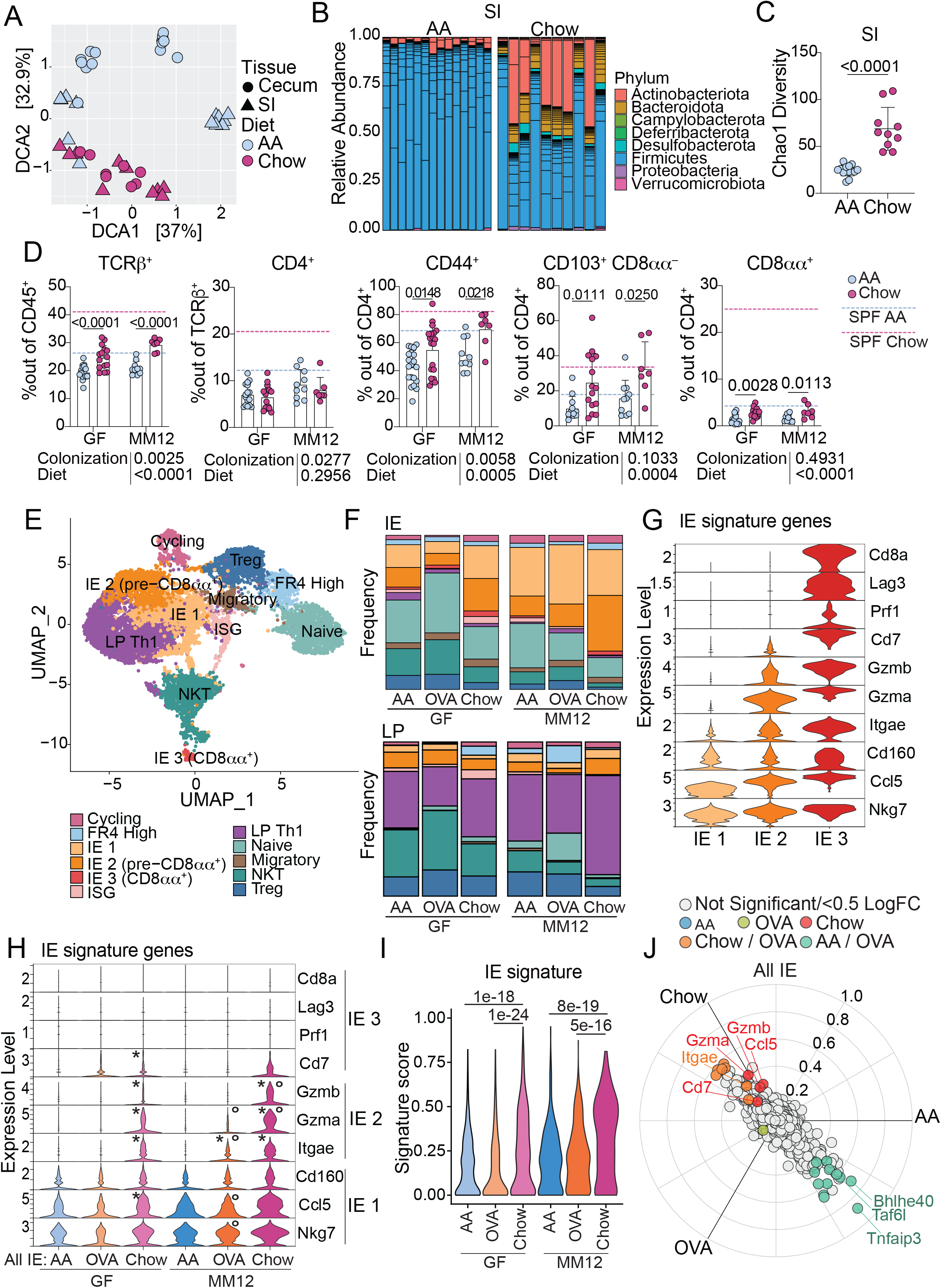
Chow diet promotes microbiota-independent epithelial adaptation and cytotoxic transcriptional programming of intestinal CD4^+^ T cells. **(A-C)** 16S rRNA sequencing of small intestine (SI) or cecum contents of 8-week-old SPF mice fed AA or standard chow diet represented by detrended correspondence analysis (DCA) (A), relative SI phyla abundance (B), and SI Chao1 alpha diversity with mean +/-SD and unpaired t-test (C). Data is from 4 independent experiments using 11-15 mice per condition. **(D)** Flow cytometry from the IE of GF or Oligo-MM^12^ mice fed AA or standard chow diet measuring frequency of the indicated cell subsets. Dashed lines show mean value from SPF Chow (red) or SPF AA (blue). Mean + SD from 3-5 independent experiments with 7-16 mice per condition. Two-way ANOVA p-values beneath each plot, and p-values <0.05 from Holm-Šidák multiple comparison test between diets within each colonization on each plot. **(E-J)** scRNAseq of 12,139 IE and LP CD4^+^ T cells from GF or Oligo-MM^12^ mice fed AA, AA+OVA, or standard chow diet with 2-4 mice per condition. **(E)** UMAP visualization of sequenced cells positioned by gene expression similarity and colored by gene expression cluster. **(F)** Frequency of cells within each cluster from the IE (top) or LP (bottom). **(G-H)** Expression (Pearson residuals) of IE signature genes within the 3 IE mature clusters (G) or within all IE CD4^+^ T cells (H). For H, Wilcoxon rank sum test with Bonferroni correction for multiple comparison, p-adj < 1e-5 were considered statistically significant. Groups labeled with asterisk (*) are significantly higher than AA diet mice within the same colonization group. Groups labeled with a circle (∘) are significantly higher than GF mice from the same dietary group. **(I)** IE gene signature score grouped by condition. Each data point contributing to the violin plots represents a single sequenced cell. Wilcoxon rank sum test with p-adj < 1e-5 within each colonization group displayed on the plot. **(J)** 3-way volcano plot showing differential gene expression between diets in all sequenced IE CD4^+^ T cells. Colored genes are differentially expressed (p-adj <0.05 from FDR-corrected Kruskal-Wallis Test and log2 fold change >0.5), colored by the diet(s) in which they are upregulated. Select genes of interest are labeled on each plot.

Similar to SPF, intestinal length was reduced in GF AA-diet mice compared to chow diet (*see* **Figure S1C),** and the tissue showed no evidence of damage, inflammation, or major morphological changes (*see* **Figure S1D-E)**. GF chow diet mice had elevated frequencies of LP macrophages, but no other differences between the diets were observed within the myeloid compartment (*see* **Figure S1F)**.

GF and Oligo-MM^12^ mice had highly reduced intestinal CD4^+^ T cells, similar to levels seen in SPF AA diet mice **(Figure 2D, Figure S2K-L)**, suggesting that both complex diet and a complex microbiota are required for steady state gut T cell accumulation. Within GF or Oligo-MM^12^ mice, chow diet once again increased IE TCRαβ^+^ T cells relative to AA diet **(Figure 2D, Figure S2K)**. Although CD4^+^ T cell frequency was not impacted, chow diet increased frequencies of CD44^+^, pre-CD8αα^+^ (CD103^+^ CD8αα^-^), and CD8αα^+^ CD4^+^ T cell subsets **(Figure 2D, Figure S2K)**, demonstrating that dietary signals promote IE CD4^+^ T cell maturation and tissue adaptation independent of the microbiota.

To address how diet impacts functional gene expression pathways of intestinal CD4^+^ T cells in greater detail, we performed single-cell RNA sequencing (scRNA-seq) of CD4^+^ T cells from the IE and LP of GF or Oligo-MM^12^ mice fed AA, AA+OVA, or chow using the Chromium 10X (10X Genomics) platform **(Figure S3A)**. Sequenced cells were assigned to 11 major unbiased clusters which we defined based on their top differentially expressed genes **(Figure 2E, Figure S3B, Table S3)**. We observed high frequencies of naïve and NKT cells, particularly in GF or AA diet mice **(Figure 2F)**. Conversely, mature cells, which formed 3 clusters in the IE and a single major cluster defined by Th1-type genes in the LP, were increasingly frequent Oligo-MM^12^ or chow diet mice **(Figure 2F, Figure S3C)**.

We assessed expression of signature genes associated with stepwise CD4^+^ T cell transcriptional adaptation to the IE (London et al., 2021), and found that our IE mature clusters represented 3 stages of signature gene acquisition **(Figure 2G, Figure S3D, Table S3)**. IE 1 cells were least adapted to the epithelium, expressing *Nkg7*, *Ccl5*, and *Cd160* which were expressed in all 3 clusters **(Figure 2G)**. IE 2 matched our pre-CD8αα^+^ population, expressing CD103 (*Itgae*), and granzymes (*Gzma*, *Gzmb*), and were enriched in chow diet relative to AA or AA+OVA regardless of colonization **(Figure 2F-G, Figure S3E)**. IE 3, representing fully IE adapted CD4^+^ CD8αα^+^ T cells, additionally expressed *Cd8a*, *Lag3*, *Prf1* (Perforin) and *Cd7*, and were rare in all sequenced groups, confirming their dependence on complex microbiota **(Figure 2F-G, Figure S3E)**. Comparing expression of IE signature genes between conditions, we found that IE1 genes (*Nkg7*, *Ccl5*, *Cd160*) were more uniformly expressed whereas IE2 genes (*Itgae, Gzma*, *Gzmb*) were expressed almost exclusively in chow diet mice from both GF and Oligo-MM^12^ **(Figure 2H**, genes significantly upregulated by diet within each colonization are indicated by *). Indeed, when we created a gene signature using all 10 IE hallmark genes, CD4^+^ T cells from chow diet mice scored higher than AA or AA+OVA diet mice in both GF and Oligo-MM^12^ **(Figure 2I)**. Unbiased three-way comparison of differential gene expression between AA, AA+OVA, or chow diet mice across GF and Oligo-MM^12^ further confirmed that IE signature genes including *Itgae* and granzymes were among the top upregulated genes in chow diet mice **(Figure 2J, Table S4)**.

Although feeding mice a single food protein (AA+OVA) did not alter frequencies of intestinal CD4^+^ T cell subsets either by flow cytometry (data not shown) or scRNAseq (*see* **Figure 2F, Figure S3C, E),** IE CD4^+^ T cells from Oligo-MM^12^ AA+OVA mice significantly upregulated *Itgae* compared to AA and had a trending increase in *Gzma* (p=0.007) **(Figure 2H)**. This difference was not seen in GF, suggesting that a low dose of single food protein is sufficient to promote some epithelial adaptation of CD4^+^ T cells in a manner that may require co-stimulation from the microbiota.

Total Treg frequency did not vary greatly between diets in GF or Oligo-MM^12^ (*see* **Figure 2F, Figure S3F)**. However, Rorγt^-^ pTregs were increased in chow diet relative to AA diet in GF and Oligo-MM^12^ **(Figure S3F)**, supporting previous reports that these cells respond to food (Kim et al., 2016). We next assessed Treg transcriptional programs within our scRNAseq dataset, identifying 8 subclusters including one that upregulated IE signature genes (*Nkg7, Gzma, Gzmb*) **(Figure S3G-I, Table S5)**. IE signature Tregs were expanded in chow diet regardless of colonization, and trended towards increase in AA+OVA mice from Oligo-MM^12^ **(Figure 3A-B)**. These cells were transcriptionally similar to previously identified pTregs on a trajectory towards CD4^+^ CD8αα^+^ differentiation (Bilate et al., 2020). In contrast, *Il10*-high Tregs were almost exclusively found in Oligo-MM^12^ (**Figure 3A, Figure S3J)** and *Rorc* expression was confined to this population among Tregs (data not shown). This population may therefore represent Rorγt^+^ pTregs known to depend on the microbiota (Sefik et al., 2015; Yang et al., 2016) while the IE signature pTregs may represent Rorγt^-^ pTregs reported to depend on diet (Kim et al., 2016). Indeed, regardless of colonization status, chow diet Tregs upregulated IE2 signature genes (*Itgae, Gzma, Gzmb*) and scored higher for total IE gene signature **(Figure 3C-D)**. Unbiased three-way comparison of differential gene expression between AA, AA+OVA, or chow diet Tregs across GF and Oligo-MM^12^ further revealed that chow diet led to upregulation of IE signature genes (*Itgae, Gzma, Gzmb, Lag3*) and core Treg suppressive function genes (*Il10, Ctla4)*, while a distinct natural Treg-associated transcriptional profile (*Gata3, Nrp1, Cd81, Klrg1*) was enriched in AA or AA+OVA diet **(Figure 3E, Table S6)**.

**Figure 3.**
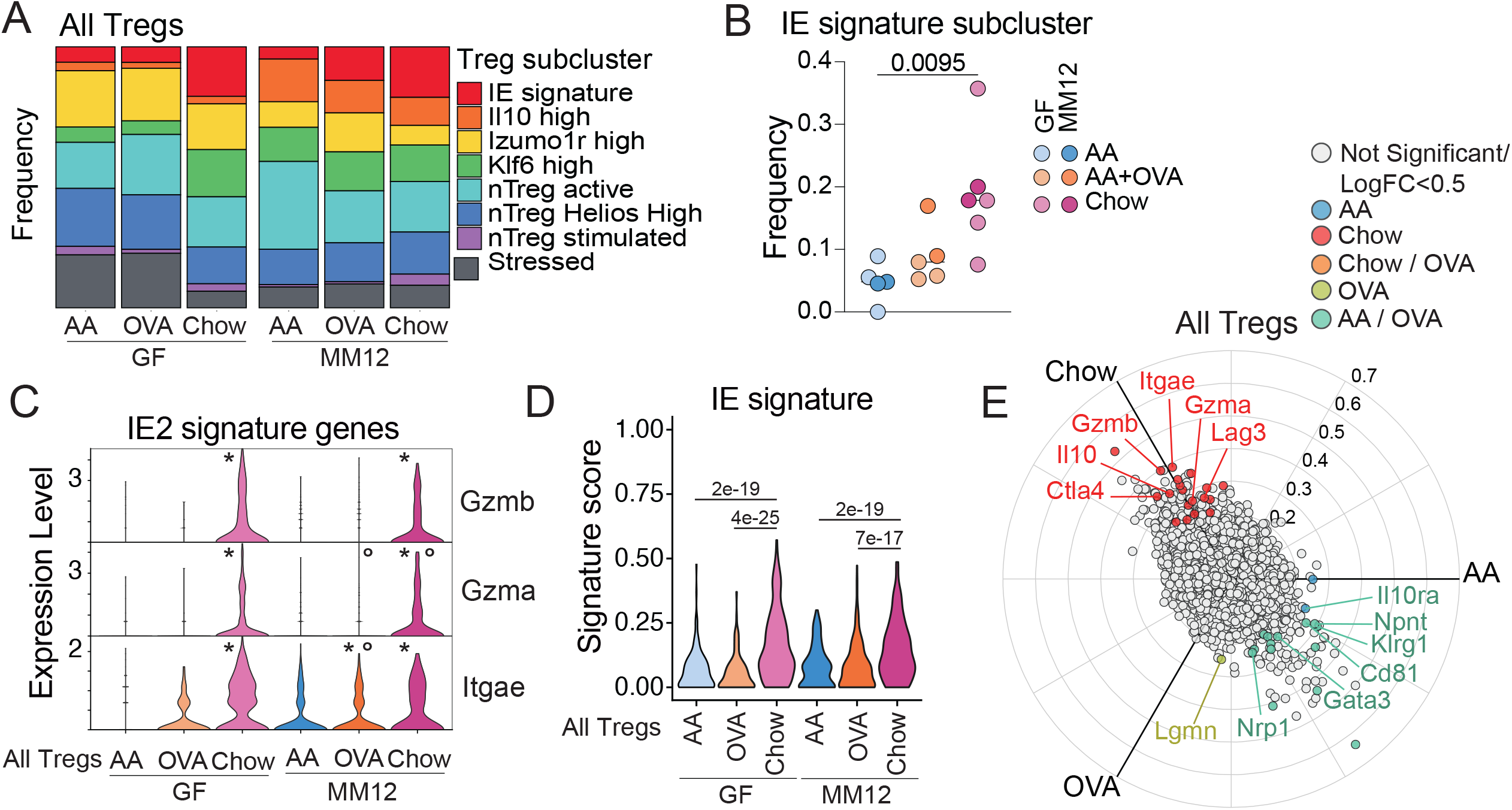
Chow diet imprints an epithelial transcriptional signature on intestinal Tregs. scRNAseq of 1183 IE and LP Tregs from GF or Oligo-MM^12^ mice fed AA, AA+OVA, or standard chow diet with 2-4 mice per condition **(A-B)** Frequency of cells in each Treg subcluster (A) or in the IE signature subcluster (B). One-way ANOVA displaying p< 0.05 from Tukey’s multiple comparisons test. **(C)** Treg expression (Pearson residuals) of IE2 signature genes, grouped by condition. Wilcoxon rank sum test with Bonferroni correction for multiple comparison, p-adj < 1e-5 were considered statistically significant. Groups labeled with asterisk (*) are significantly higher than AA diet mice within the same colonization group. Groups labeled with a circle (∘) are significantly higher than GF mice from the same dietary group. **(D)** Treg IE gene signature score grouped by condition. Each data point contributing to the violin plots represents a single sequenced cell. Wilcoxon rank sum test with p-adj < 1e-5 within each colonization group displayed on the plot. **(E)** 3-way volcano plot showing differential gene expression between diets in all Tregs. Colored genes are differentially expressed (p-adj <0.05 from FDR-corrected Kruskal-Wallis Test and log2 fold change >0.5), colored by the diet(s) in which they are upregulated. Select genes of interest are labeled on each plot.

Similar to our findings in total IE CD4^+^ T cells, a low dose of OVA did not impact Treg transcriptional profile in GF mice but was sufficient to promote *Itgae* expression in Oligo-MM^12^ **(Figure 3C)**. Correspondingly, in both total CD4^+^ T cells and Tregs, Oligo-MM^12^ promoted expression of IE signature genes including *Gzma* and *Itgae* over GF mice with the same diet **(Figure 2H**, **Figure 3C;** indicated by ∘), but this depended on the presence of food protein. Exposure to food protein therefore recruits antigen-experienced CD4^+^ T cells to the intestinal epithelium, imprinting a tissue-resident cytotoxic gene expression signature in a manner that is amplified by microbial co-stimulation.

### Complex diet induces Granzyme B expression in intestinal T cells

To further validate our findings from scRNAseq, we assessed intestinal T cell Granzyme B expression by flow cytometry. Indeed, chow diet promoted Granzyme B expression among total IE CD4^+^ T cells and Tregs in both GF and SPF **(Figure 4A)**. Additionally, Granzyme B expression was higher among CD8αβ^+^ T cells, and thymic TCRγδ^+^ and CD8αα^+^ TCRαβ^+^ T cells **(Figure 4B)**. Whereas diet-induced Granzyme B expression among CD4^+^ and CD8αβ^+^ T cells was boosted in the presence of microbiota, the impact on thymic-derived T cell subsets was microbiota-independent. A complex diet therefore broadly induces Granzyme B expression in intestinal T cells, although the pathway of induction may be distinct depending on the subset.

**Figure 4.**
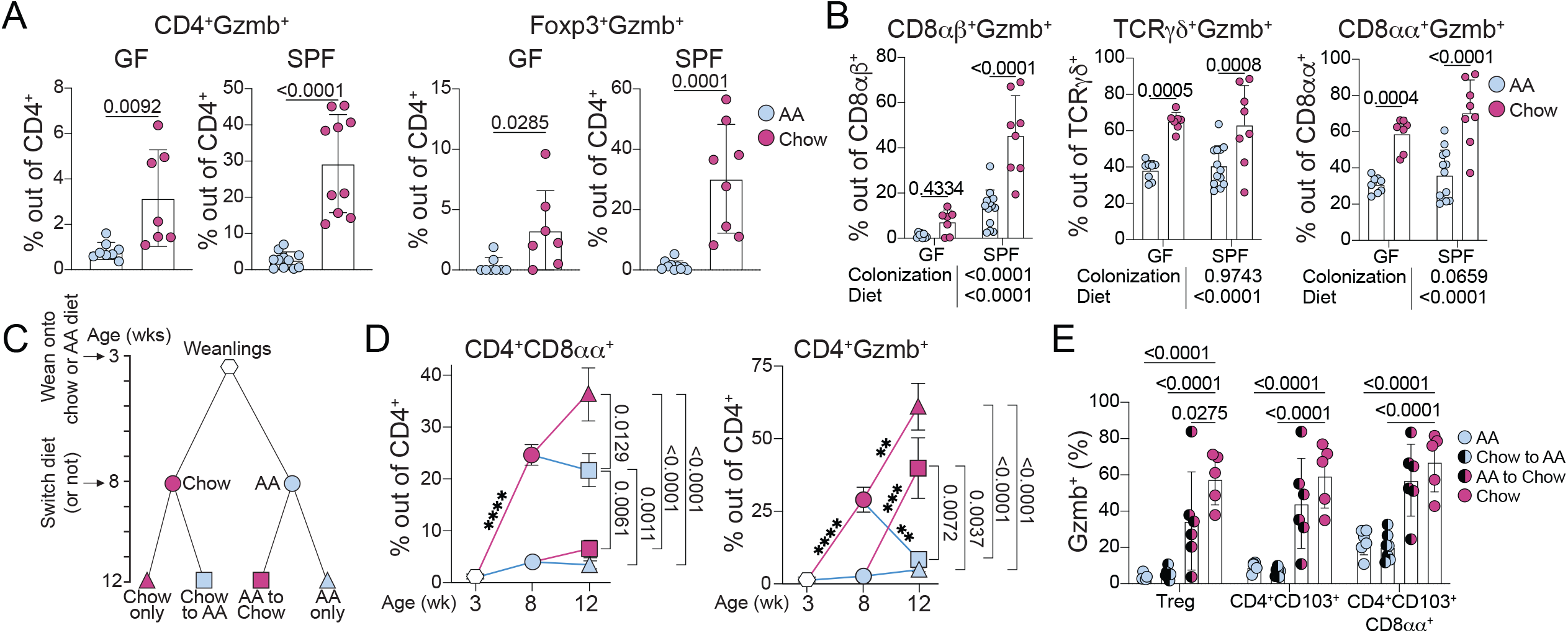
Chow diet induces Granzyme B expression in intestinal T cells. **(A-B)** Flow cytometry analysis of Granzyme B expression within IE T cell subsets from 8-week-old SPF or GF mice fed AA or standard chow diet. Mean +/-SD from 2-3 independent experiments with 7-11 mice per condition. Unpaired t-tests displaying p<0.05 (A) or two-way ANOVA p-values beneath each plot, and p<0.05 from Holm-Šidák multiple comparison test between diets within each colonization displayed on each plot (B). **(C)** Schematic of weaning and diet switch experiments. **(D)** Flow cytometry analysis of 3, 8, and 12-week-old SPF mice fed chow or AA diets according to schematic C. Mean +/-SEM from 2-5 independent experiments with 5-18 mice per condition. One-way ANOVA comparing conditions in 12-week-old mice displaying p<0.05 from Tukey’s multiple comparison test to the right of each plot. Unpaired t-tests comparing consecutive timepoints between conditions with Holm-Šidák correction for multiple comparisons displayed on plot, **p < 0.01, ***p<0.001, ***p<0.0001. **(E)** Flow cytometry of Granzyme B expression within IE T cell subsets from 12-week-old SPF mice fed chow or AA diets according to shematic C. Mean +/-SD from 2 independent experiments with 5-7 mice per condition. Two-way ANOVA displaying p<0.05 from Dunnett’s multiple comparison test comparing each group against chow only.

To characterize the kinetics of dietary imprinting on intestinal CD4^+^ T cells, SPF mice were analyzed at 3 weeks (pre-weaning), 8 weeks, or 12 weeks old. Some mice that were fed AA or chow diet until 8 weeks old (*ie*. following the standard protocol for prior experiments) were switched to the opposite diet until endpoint analysis at 12 weeks old **(Figure 4C)**. This experimental setup enabled us to assess intestinal CD4^+^ T cell fate if dietary signals were removed from adult mice (Chow to AA) or if exposure to complex diet was delayed until adulthood (AA to Chow). Pre-weaning (3-week-old) mice had very low frequencies of CD8αα^+^ or Granzyme B+ CD4^+^ T cells **(Figure 4D)**. Mice that were subsequently weaned onto chow diet had progressively increased frequencies of these cells by 8 and then 12 weeks old, whereas AA diet mice maintained a weanling-like immature IE CD4^+^ T cell profile **(Figure 4D)**. Switching AA to chow diet at 8 weeks did not recover CD4^+^ CD8αα^+^ T cell frequency, but did rescue Granzyme B expression **(Figure 4D**, red squares). Conversely, chow-induced CD4^+^ CD8αα^+^ T cells persisted in mice switched to AA diet at 8 weeks whereas Granzyme B expression declined **(Figure 4D**, blue squares). This may indicate a critical window during weaning and development during which diet imprints an epithelial signature on intestinal CD4^+^ T cells, whereas once CD4^+^ CD8αα^+^ develop they can persist at least 4 weeks without chow diet. By contrast, Granzyme B expression is dependent on recent exposure to a complex diet.

Our previous work has demonstrated that IE adaptation by CD4^+^ T cells is linked to Granzyme B upregulation (Bilate et al., 2020; London et al., 2021; Mucida et al., 2013; Reis et al., 2013). However, in these diet switch experiments, Granzyme B expression and IE adaptation were uncoupled such that Chow-to-AA mice have IE-adapting CD4^+^ T cells that no longer express Granzyme B **(Figure 4E)**. Altogether, these data demonstrate that dietary signals in conjunction with the microbiota promote both CD4^+^ CD8αα^+^ differentiation and Granzyme B expression in intestinal CD4^+^ T cells though the temporal dynamics and therefore pathways of induction appear distinct.

### Exposure to dietary protein drives clonal selection of intestinal CD4^+^ T cells

We next assessed how routine exposure to dietary protein shapes the TCR repertoire of intestinal CD4^+^ T cells by analyzing TCRs from our scRNAseq of GF and Oligo-MM^12^ mice fed AA, AA+OVA, or chow. IE and LP mature CD4^+^ T cells were overall highly clonally expanded despite the absence of complex microbiota, whereas naïve cells and NKT cells (which use an invariant TCRα but diverse TCRβ) were highly diverse as expected **(Figure 5A)**. We estimated repertoire diversity across sample groups using D50 in which repertoires are scored from 0 (least diverse) to 0.5 (most diverse) and found a trend towards higher clonal diversity in GF AA or AA+OVA compared to the other groups (**Figure 5B)**. However, within mature (IE1, IE2, IE3, LP Th1) CD4^+^ T cells or Tregs we did not observe major differences in repertoire diversity across sample groups (**Figure 5B),** demonstrating that while few mature CD4^+^ T cells accumulate in the intestine in absence of major foreign antigen exposure, those that do are still clonally expanded. We recently reported that gut-associated germinal centers are highly reduced in GF AA diet mice, although B cells from these germinal centers still exhibit clonal selection and expansion (Nowosad et al., 2020), suggesting a parallel mechanism for intestinal B cells.

**Figure 5.**
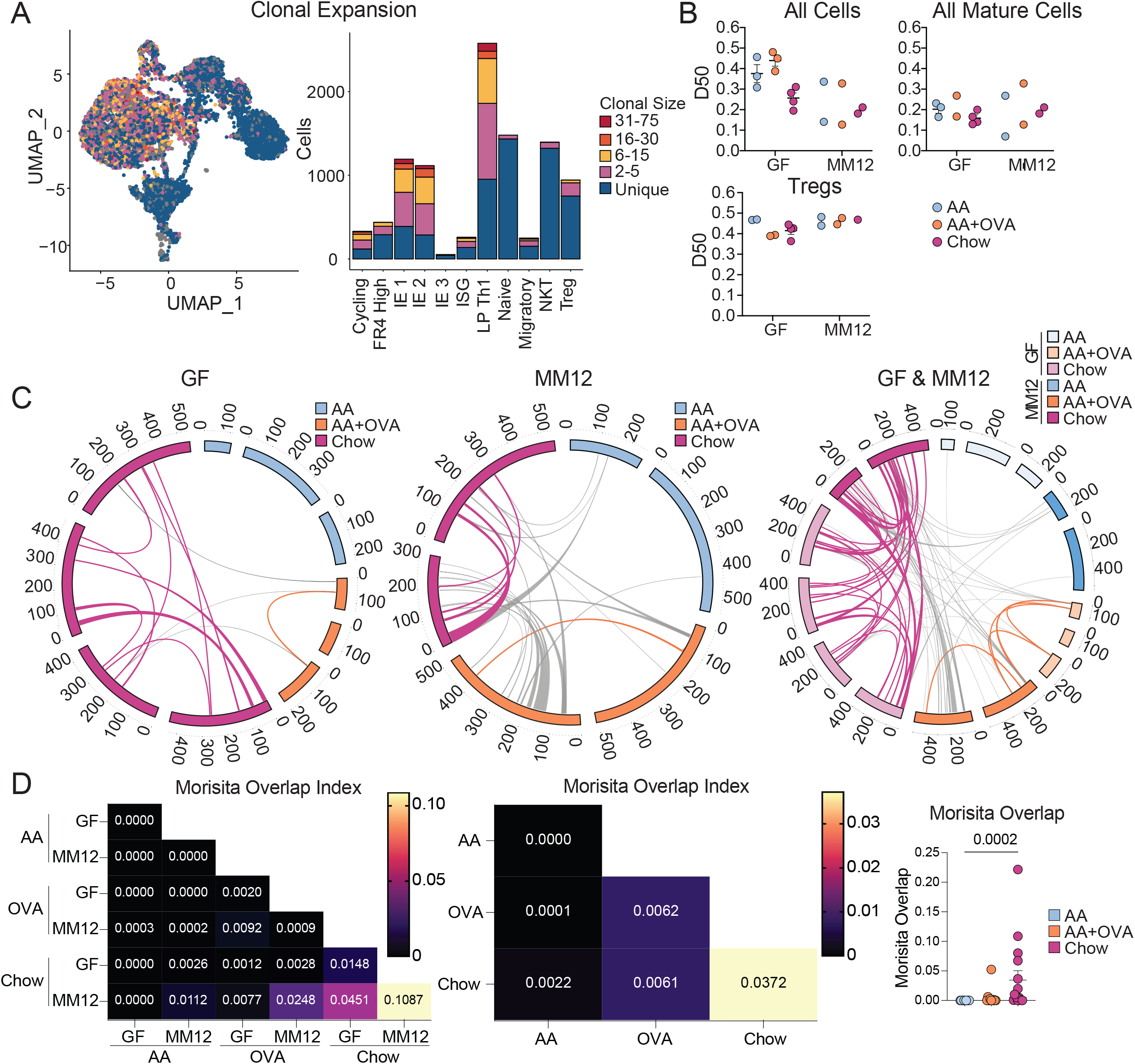
Exposure to dietary protein drives clonal selection of intestinal CD4^+^ T cells. scTCRseq of 12,139 IE and LP CD4^+^ T cells from GF or Oligo-MM^12^ mice fed AA, AA+OVA, or standard chow diet using 2-4 mice per condition. **(A)** Clonal expansion (by TCR nucleotide sequence) of cells visualized by UMAP (left) and barplot of gene expression clusters (right). **(B)** D50 in which repertoires are scored from 0 (least diverse) to 0.5 (most diverse) within all cells (top left), mature clusters IE1, IE2, IE3, and LP Th1 combined (top right), and Tregs (bottom left). **(C-D)** Clonal sharing between mice defined by paired TCRα and TCRβ CDR3 amino acid sequence. NKT cells were discarded from analysis. **(C)** Circos plots in which each segment represents a mouse, colored by diet and sized by cell count. Links between segments represent public clones which are colored by diet if shared between mice of the same diet or uncolored if shared between mice of different diets. **(D)** Morisita overlap index heatmaps where each square represents the mean overlap between each mouse in the indicated conditions (left and center) or scatter plot where each dot represents overlap between mice in the same diet (right). Kruskal-Wallis test with p<0.05 from Dunn’s multiple comparisons displayed on plot.

To determine whether food antigens can drive clonal selection of intestinal CD4^+^ T cells, we examined shared (public) clones between mice, as defined by identical paired TCRα and TCRβ CDR3 amino acid sequences (**Figure 5C-D)**. Among GF mice where the primary source of foreign antigen is food, we found no clonal overlap between AA diet mice, a low level of clonal sharing between AA+OVA mice, and extensive clonal sharing between chow diet mice, raising the possibility that food antigen is driving clonal selection. We additionally found a small amount of clonal overlap between AA+OVA and chow diet mice, which may be driven by self or unaccounted environmental antigens. There was more clonal overlap in general between Oligo-MM^12^ mice, likely due to recognition of microbial antigen and/or a co-stimulatory effect from microbiota. Nevertheless, between diets in Oligo-MM^12^ we found similar trends to GF, with no clonal sharing between AA diet mice, low level sharing between AA+OVA mice, and a higher degree of sharing between chow diet mice. When we compared clonal overlap across GF and Oligo-MM^12^ mice, we found some additional sharing between AA+OVA mice, whereas clones were highly shared between chow diet mice. These data point to clonal selection of intestinal CD4^+^ T cells by dietary antigen, where exposure to a low dose of a single food protein leads to a low level of clonal selection, while exposure to diverse food proteins in the context of complex chow results in a high level of selection. In this case, presence of diverse dietary metabolites may have an additional adjuvant effect, promoting differentiation and expansion of food antigen-specific T cells.

### Context of exposure shapes intestinal CD4^+^ T cell responses to food protein

To better understand how food-induced gut CD4^+^ T cells contribute to tolerance, and how these responses are perturbed in the context of food allergy, we combined a naïve T cell fate-mapping mouse model (iSell^Tomato^) (Merkenschlager et al., 2021) with a cholera toxin (CT) mouse model of food allergy (Jimenez-Saiz et al., 2017). iSell^Tomato^, in which CD62L^+^ naïve cells are permanently labeled with Tomato fluorescence upon tamoxifen administration, enables identification of “ex-naïve” (Tomato^+^ CD62L^−^) T cells that matured and entered the gut since the time of tamoxifen labeling (Parsa et al., 2022). We utilized this strategy to enrich for intestinal CD4^+^ T cells responding to dietary protein in 3 contexts: 1) Feeding, where mice are fed OVA, representing the steady state response to food; 2) Allergy, where primary exposure to OVA is with CT, resulting in allergic sensitization; 3) Tolerance, where mice are fed OVA alone prior to OVA/CT, resulting in oral tolerance and protection against allergy **(Figure 6A)**. Tomato^+^ CD62L^−^ cells analyzed at day 26 of our treatment protocol therefore include all cells that matured and entered the gut during the 4-week treatment protocol, which will be enriched for cells responding to OVA but may also include cells responding to CT, microbiota, and other food antigens, since CD4^+^ T cells are continuously recruited to the intestine. At the day 26 timepoint, mice in the allergy group have elevated serum total IgE and OVA-specific IgG1, demonstrating allergic sensitization **(Figure 6B),** but have no observable increase in intestinal tissue damage or inflammation **(Figure S4A)**. Upon continuation of this protocol for 4 weekly doses of OVA/CT, systemic OVA challenge results in anaphylaxis in allergy mice (Jimenez-Saiz et al., 2017), while mice in the tolerance group are protected **(Figure 6C)**.

**Figure 6.**
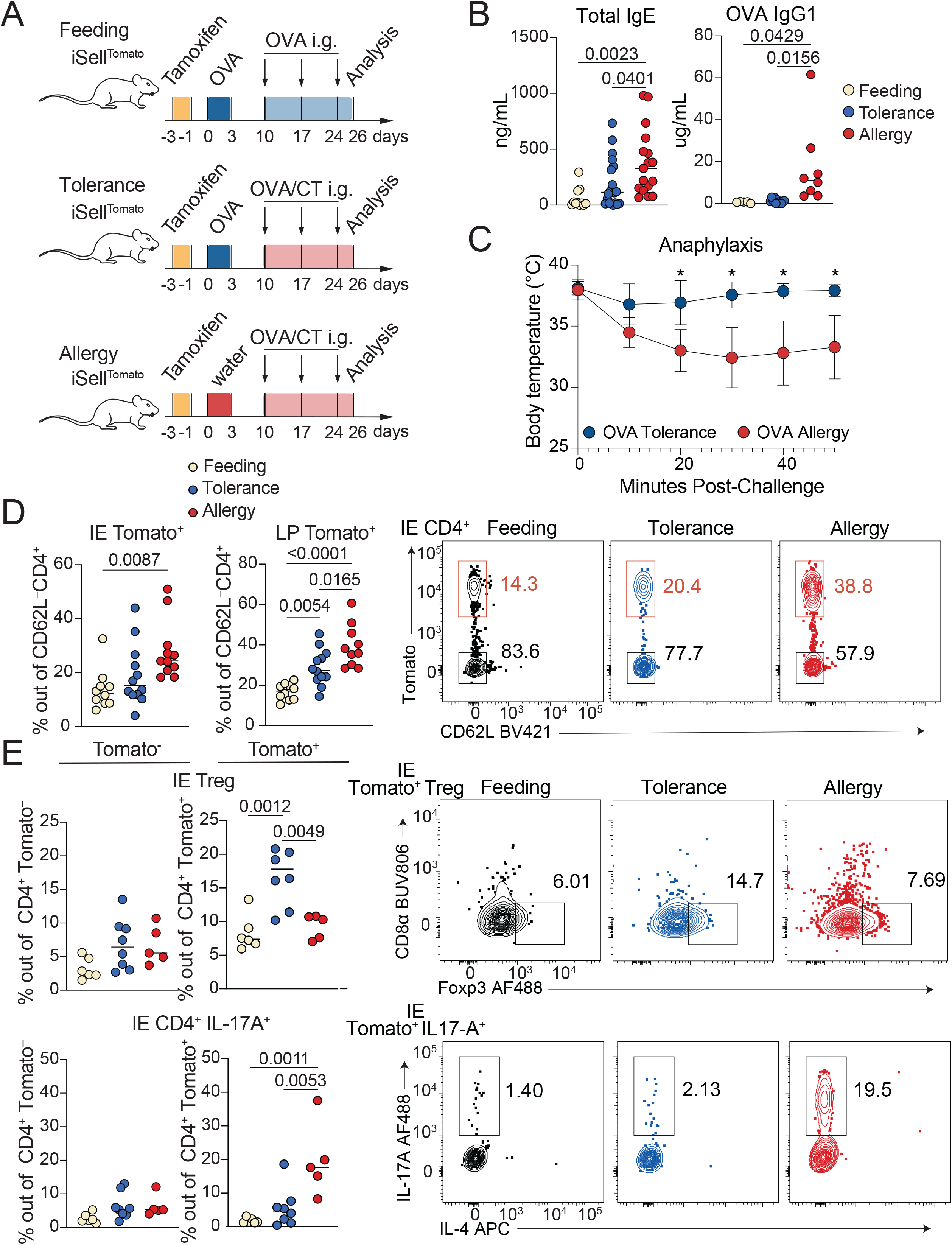
Tracking intestinal CD4^+^ T cell responses during OVA feeding, tolerance, or allergy. iSell^Tomato^ mice were analyzed on day 26 after treatment with tamoxifen to permanently label naïve T cells and then exposure to OVA in the context of feeding, tolerance, or allergy. **(A)** Schematic of experimental protocol. **(B)** Total serum IgE (left) or OVA specific IgG1 (right) as measured by ELISA. One-way ANOVA displaying p<0.05 from Tukey multiple comparison test. Data is representative of 2-3 independent experiments with 4-13 mice per group. **(C)** Anaphylaxis as measured by body temperature of mice at the indicated times after intraperitoneal OVA injection, following 4 weekly doses of OVA/CT. Mean +/-SEM representative of 2 independent experiments with 5-6 mice per group. Unpaired t tests with Holm-Sidak multiple comparison test, *p<0.05. **(D-E)** Flow cytometry measuring frequency of Tomato^+^ out of total CD4^+^ T cells in the IE or LP (D) or of the indicated CD4^+^ T cell subsets out of Tomato^+^ or Tomato^-^ CD4^+^ T cells in the IE (E) with representative flow cytometry plots shown on the right. Mean from 4 (D) or 2 (E) independent experiments with 10-12 (D) or 5-8 (E) mice per group. One-way ANOVA with Tukey multiple comparison test, showing p-values <0.05.

Four weeks after initial tamoxifen administration, ∼15% of small intestine IE or LP CD4^+^ T cells from SPF mice in the OVA feeding protocol were Tomato^+^ CD62L^−^ **(Figure 6D)**, representing the baseline for CD4^+^ T cell influx to the intestine at steady state. In both tissues, CD4^+^ T cell influx was increased over 2-fold in OVA allergy mice, while OVA tolerance fell between feeding and allergy **(Figure 6D)**. By contrast, in the large intestine there was no increase in CD4^+^ T cell influx in tolerance or allergy mice **(Figure S4B)**, suggesting that responses to food in inflammatory contexts remain localized to the small intestine. Among Tomato^−^ (pre-existing non-naïve) CD4^+^ T cells, mice from each treatment group had similar frequencies of IE2, IE3, Tregs, and cells producing IL-17A or IL-4 based on flow cytometry analysis (**Figure 6E, Figure S4C-D)**. However, profiles of infiltrating Tomato^+^ CD4^+^ T cells were distinct in OVA feeding, tolerance, and allergy conditions, suggesting that iSell^Tomato^ is an effective system for enriching and characterizing polyclonal intestinal CD4^+^ T cells responding to food protein in different contexts. In OVA tolerance, Treg frequency was higher among Tomato^+^ CD4^+^ T cells whereas in OVA allergy there was increased IL-17A expression **(Figure 6E, Figure S4D)**, consistent with prior reports that CT induces intestinal Th17 in a microbiota-dependent manner (Zhao et al., 2017). C57BL/6 mice do not typically have a strong Th2 response in the CT allergy model and we did not see significant increases in IL-4 production among Tomato^+^ CD4^+^ T cells from allergy mice **(Figure S4C-D)**. IE Tomato^+^ CD4^+^ T cells from all three treatment groups demonstrated only partial acquisition of an epithelial profile in the course of four weeks, with ∼25% expressing CD103 (IE2) but few cells expressing CD8αα (IE3) **(Figure S4C)**.

To transcriptionally characterize the CD4^+^ T cell response to OVA in the context of feeding, tolerance, and allergy, we performed scRNA-seq on Tomato^+^ CD62L^−^ (fate-mapped ex-naïve) and Tomato^−^ CD62L^−^ (pre-existing non-naïve) CD4^+^ T cells (pooled at a ratio of ∼1:2 Tomato^+^:Tomato^−^ to evenly enrich for Tomato^+^ cells) from the IE and LP of SPF iSell^Tomato^ mice on day 26 of the treatment protocols (*see* **Figure 6A)**. scRNA-seq was performed on the 10X Genomics platform with 4-5 mice per condition pooled across 2 independent experiments and sequencing runs **(Figure S4E-F)**. We expect that our designated Tomato^-^ and Tomato^+^ populations represent enrichment for true pre-existing mature versus ex-naïve cells, whereas there may be some contaminating cells in each population due to incomplete efficiency of the fate mapping mouse model, low detection of *TdTomato/Stop*, or labeling of Sell^+^ central memory cells which could potentially differentiate and enter the gut.

Sequenced cells were assigned to 11 major unbiased clusters which we defined based on their top differentially expressed genes **(Figure 7A, Figure S4G-H, Table S7)**. Tomato^+^ cells were present in all clusters, though they were particularly enriched among CCR7+ migratory cells and Tregs and were lowest in IE-adapted clusters (discussed below) **(Figure S4F)**. Consistent with a weak Th2 response in the C57BL/6 CT allergy model, we did not identify gene expression clusters defined by hallmark Th2 genes **(Figure S5A)**. Based on expression of IE signature genes, we identified IE2 and IE3 clusters comparable to our GF/Oligo-MM^12^ dataset, but no IE1 cluster, further reinforcing the notion of an additive effect of chow diet and full colonization on CD4^+^ T cell IE adaptation **(Figure 7A, Figure S5B, Table S7)**. An additional IE cluster was identified (IE4), which expressed the full IE3 signature as well as additional genes associated with NK function (*Fcer1g*, *Tyrobp*, *Klrd1*), Treg function (*Ikzf2*), and apoptosis (*Tox*, *Bcl2*), while lower *Cd4* levels **(Figure S5B, Table S7)**. Tomato labeling frequency progressively decreased from IE2 (40%) to IE3 (15%) to IE4 (5%), confirming that only partial acquisition of an epithelial profile occurs in the course of four weeks **(Figure S5C)**. Comparison of cluster distributions between groups demonstrated that IE4 cells were almost exclusively Tomato^+^ and from either tolerance or allergy mice **(Figure S5D, Figure 7B)**. Indeed, three-way differential gene expression analysis among IE Tomato^-^ cells showed that IE4 genes were highly enriched in tolerance and allergy groups compared to feeding **(Figure S5E, Table S8)**. These data suggest that the IE4 gene expression program was turned on in pre-existing tissue resident IE3 cells in inflammatory (CT-exposed) conditions. IE4 may therefore represent an activated state of IE-adapted CD4^+^ T cells in which they can function in an antigen-independent manner through innate receptors, as previously suggested (Bilate et al., 2020). Of note, the IE4 transcriptional program is similar to thymic-derived (CD4^-^) CD8αα^+^ TCRαβ^+^ cells found in both tumors and the steady state intestinal epithelium (Chou et al., 2022; Denning et al., 2007), though with the addition of *Ikzf2* which may suggest some regulatory capacity.

**Figure 7.**
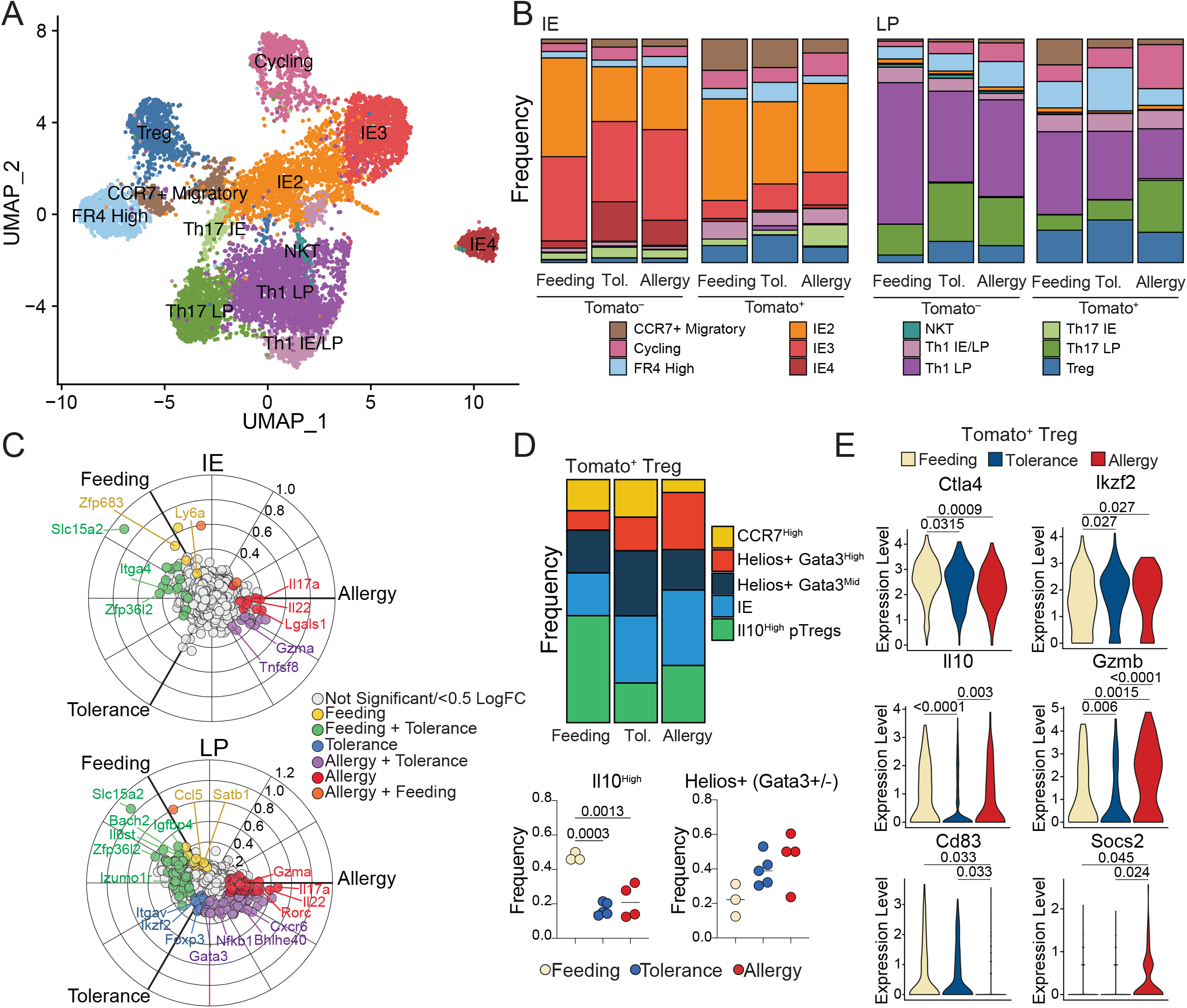
Distinct intestinal CD4+ T cell responses to OVA feeding, tolerance, and allergy. scRNAseq of 11,217 Tomato+ and Tomato-CD4+ T cells from the IE and LP of mice on day 26 of OVA feeding, tolerance, or allergy protocols using 4-5 mice per condition pooled across 2 independent experiments. **(A)** UMAP visualization of sequenced cells positioned by gene expression similarity and colored by gene expression cluster. **(B)** Frequency of cells within each cluster from the IE (left) or LP (right) within each sample group. **(C)** 3-way volcano plots showing differential gene expression between conditions in Tomato+ CD4+ T cells from the IE (top) or LP (bottom). Colored genes are differentially expressed (p-adj <0.05 from FDR-corrected Kruskal-Wallis Test and log2 fold change >0.5), colored by the condition(s) in which they are upregulated. Select genes of interest are labeled on each plot. **(D)** Analysis of Treg subclusters among Tomato+ Tregs (496 total cells) showing frequency of all subclusters (above) or Il10+ or pooled Helios+ subsets displaying p<0.05 one-way ANOVA with Tukey multiple comparison test (below). **(E)** Differentially expressed Treg functional genes between Tomato+ Tregs (496 total cells) in different conditions displaying p-adj <0.05 from Wilcoxon rank sum test corrected with FDR.

To assess whether CD4^+^ T cells that mature and enter the gut during OVA feeding, tolerance, or allergy have distinct characteristics, we compared cluster distributions of Tomato^+^ cells between these groups. In agreement with our observations from flow cytometry, OVA tolerance mice had increased frequencies of Tregs whereas allergy mice had increased Th17 among IE Tomato^+^ cells; a similar though non-significant trend was observed in the LP **(Figure 7B, Figure S5F)**. Three-way differential gene expression found that IE and LP Tomato^+^ cells from OVA feeding mice upregulated genes associated with tissue residency and homing (*Zfp683, Ccl5*), memory (*Ccr7*, *Satb1*, *Bach2*) and type I immune responses (*Ly6a, Ifi47*). By contrast, OVA allergy Tomato^+^ cells upregulated Th17-associated genes (*Il17a, Il22, Rorc, Them176a*) **(Figure 7C, Table S9-10)**. OVA tolerance mice shared the majority of their upregulated genes with feeding or allergy mice demonstrating their intermediate inflammatory phenotype. However, in the LP, tolerance alone upregulated Treg associated genes (*Foxp3, Ikzf2, Itgav, Cd83*) **(Figure 7C)**. Finally, LP Tomato^+^ cells from both tolerance and allergy upregulated *Gata3,* which was primarily expressed in Tregs in this sequencing dataset **(Figure 7C, Figure S5A)**.

We next assessed Treg transcriptional programs in detail, identifying 5 subclusters based on top differentially expressed genes **(Figure S5G, Table S11)**. Among Tomato^+^ Tregs, subcluster frequencies varied between conditions. At steady state (OVA feeding), ∼50% of incoming Tregs were *Il10* high pTregs, whereas this dropped to ∼20% in either OVA tolerance or allergy conditions **(Figure 7D)**. Incoming Tregs in these settings instead trended for enrichment of Helios^+^ (*Ikzf2*) cells (feeding vs. allergy p = 0.059), which were primarily either *Gata3*-high in allergy or *Gata3*-intermediate in tolerance **(Figure 7D)**. Tomato^+^ Tregs also differentially expressed key functional molecules between conditions **(Figure 7E)**. Compared to steady state, both tolerance and allergy conditions suppressed *Ctla4* and upregulated *Ikzf2*. Tolerance Tomato^+^ Tregs downregulated *Il10 and Gzmb* relative to the other two conditions whereas allergy promoted *Gzmb.* Allergy was additionally characterized by decreased *Cd83* and increased *Socs2*. Tregs that infiltrate the intestine during active tolerance or allergy therefore bear distinct transcriptional profiles which may contribute to the respective functional outcomes.

Altogether, these findings demonstrate how steady state CD4^+^ T cell gut infiltration and tissue adaptation, imprinted by signals from both diet and the microbiota, is disrupted by an inflammatory challenge resulting in an effector CD4^+^ T cell response. Maintenance of immune tolerance to food protein during inflammatory challenge is associated with increased influx of Tregs, whereas in the OVA/CT allergy model we observed increased influx of pro-inflammatory Th17. In both tolerance and allergy, we found altered functional gene expression programs in incoming Tregs, as well as altered transcriptional programs in resident IE CD4^+^ T cells. These data demonstrate that distinct intestinal immune responses underly steady state exposure to food versus active tolerance to food upon inflammatory challenge.

### Clonal dynamics and antigen-specificity of tolerogenic and inflammatory CD4^+^ T cell responses to food

To gain insight into the polyclonal repertoire dynamics of CD4^+^ T cell responses to food protein in the context of feeding, tolerance, or allergy, we next assessed TCRs from our scRNAseq dataset. Under steady state conditions, we found a high degree of clonal expansion among most clusters, although Tregs and FR4 high cells were more clonally diverse **(Figure 8A)**. The TCR repertoires of Tomato^-^ cells were equally diverse across conditions, whereas Tomato^+^ cells trended towards increased clonal expansion in OVA tolerance or allergy **(Figure 8A-B)**. Finally, we found increased clonal expansion of Tregs in the tolerance group, and a trend towards increased expansion of Th17 cells in the allergy group (p=0.20, **Figure 8C)**, suggesting that these subsets are simultaneously recruited and expanded in each respective condition.

**Figure 8.**
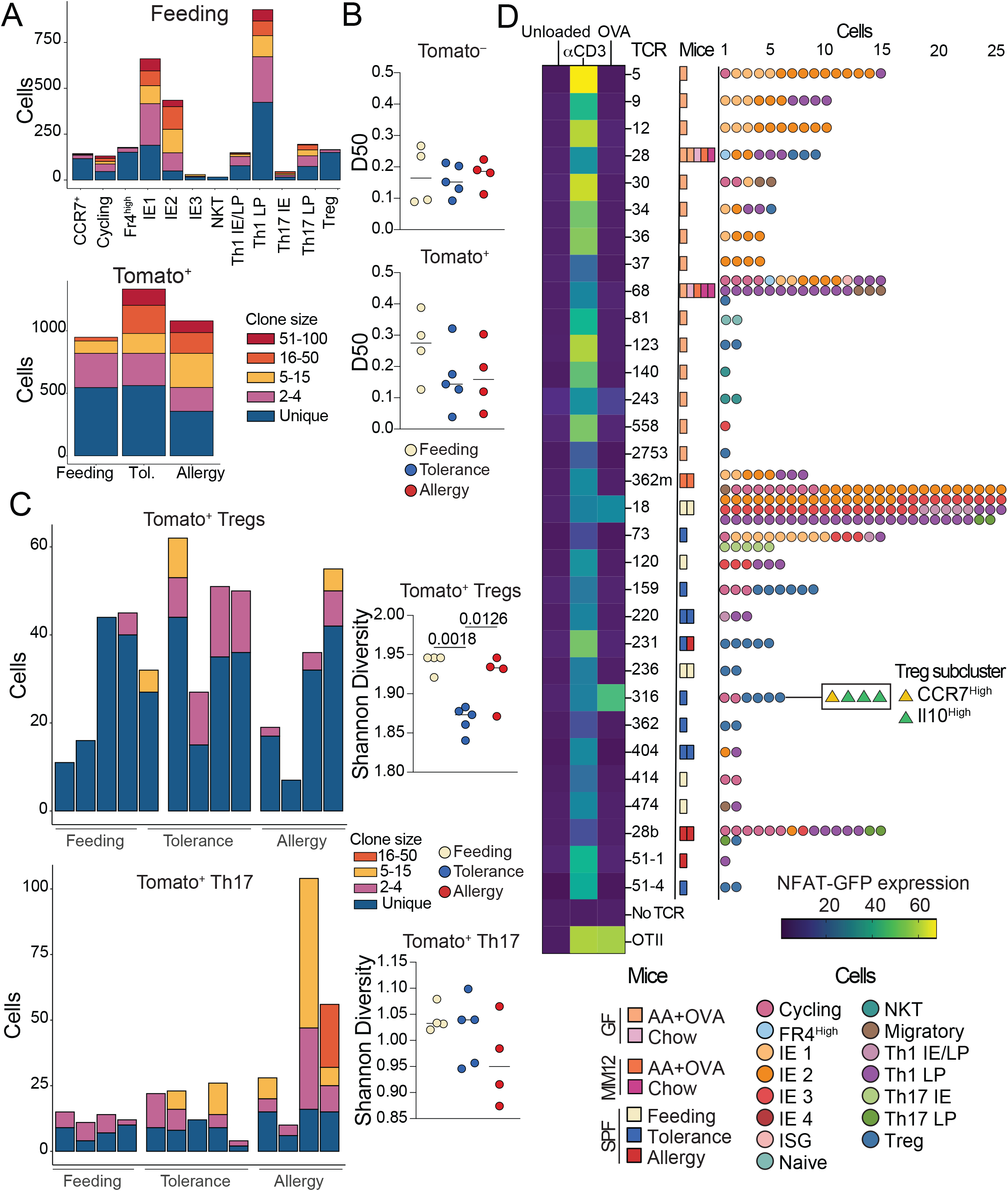
Clonal dynamics and antigen-specificity of tolerogenic and inflammatory CD4+ T cell responses to food. **(A-C)** scTCRseq of 11,217 Tomato+ and Tomato-CD4+ T cells from the IE and LP of mice on day 26 of OVA feeding, tolerance, or allergy protocols using 4-5 mice per condition pooled across 2 independent experiments. **(A)** Clonal expansion size (by TCR nucleotide sequence) plotted by gene expression cluster within all OVA feeding cells (top), and by mouse within all Tomato+ cells (bottom). **(B)** D50 in which repertoires are scored from 0 (least diverse) to 0.5 (completely diverse) within Tomato- or Tomato+ cells from each mouse. **(C)** Clonal expansion size (by TCR nucleotide sequence) among Tomato+ Tregs (above) or Th17 (below) with corresponding Shannon diversity scores to the right. One-way ANOVA with Tukey multiple comparison test, showing p-values <0.05 (B, C). **(D)** NFAT-GFP assay to determine TCR recognition of OVA relative to a-CD3 (positive control) or unloaded DCs (negative control). Heatmap indicates percent NFAT-GFP expression out of TCR+ NFAT hybridoma cells. Mouse experimental group and scRNAseq cluster of cells from which each TCR was identified are indicated to the right.

Finally, we assessed whether TCRs identified in our scRNAseq datasets were specific for dietary protein. We selected 31 TCRs from GF mice fed AA+OVA diet (*ie.* where OVA is the only source of foreign antigen) or from Tomato^+^ cells in OVA feeding or tolerance conditions **(Table S12)**, and expressed them in NFAT-GFP hybridomas (Ise et al., 2010). Two TCRs responded to OVA in the NFAT-GFP assay – one from OVA feeding (TCR 18) and one from OVA tolerance (TCR 316) **(Figure 8D)**. We further tested these two TCRs for reactivity in an overlapping OVA peptide library and found that they both bind within the epitope OVA 26:40 (**Figure S5H)**. By contrast, the OT-II TCR, which was generated in response to OVA through immunization, binds OVA 323:339 **(Figure S5H)**. This may suggest that T cell responses generated upon oral exposure to food proteins preferentially use different epitopes than immunizing responses.

TCR 18, which derived from a Tomato^+^ OVA feeding mouse, was highly expanded and found primarily within IE2, IE3 and Th1 clusters **(Figure 8D)**. TCR 316, derived from Tomato^+^ OVA tolerance, was moderately expanded and found only within Treg and cycling clusters **(Figure 8D)**. Further, these Tregs were primarily from the *Il10*+ subcluster, suggesting that although infiltration of *Il10*+ Tregs decreased in tolerance compared to steady state, these cells still have an antigen-specific functional role in maintaining oral tolerance. These findings demonstrate that in a fully polyclonal system, food antigen-specific T cell responses are generated not only in active oral tolerance, but also in the course of natural feeding.

## Discussion

Here we provide a comprehensive analysis of intestinal CD4^+^ T cell clonal dynamics and functional differentiation in response to food. We found that at steady state, signals from food promote clonal selection, epithelial adaptation, and cytotoxic programming of intestinal CD4^+^ T cells, a pathway which is further boosted by signals from the microbiota. The requirement for Tregs in controlling inflammatory responses to food is well established (Mucida et al., 2005; Torgerson et al., 2007) and an LP Treg response to dietary protein has been described using TCR monoclonal models (Hadis et al., 2011), MHC Class II tetramers (Hong et al., 2022), and antigen-free dietary models (Kim et al., 2016). Our data further demonstrates that diet-induced Tregs in the gut upregulate genes associated with cytotoxicity and epithelial residency at steady state. Previously, we reported that IE Tregs can differentiate into CD4^+^ CD8αα^+^ T cells in a microbiota dependent manner (Sujino et al., 2016), and that Tregs expressing IE signature genes are likely on a trajectory towards this fate (Bilate et al., 2020). These data suggest that dietary signals in the intestine promote a cytotoxic IE-adapted phenotype not only in conventional CD4^+^ T cells but also in Tregs.

While steady state dietary signals promoted Granzyme B expression in Tregs, we showed that incoming Tregs in the allergy model further upregulated Granzyme B, whereas tolerant mice suppressed it, indicating that regulation of this pathway may be important for Treg-mediated immune control. Granzyme-dependent cytolysis of activated effector cells has been described as one mechanism of Treg suppressive function (Grossman et al., 2004), suggesting that cytotoxicity can contribute to both pro- and anti-inflammatory responses. CD4^+^ T cells with cytotoxic properties have also been described in the setting of chronic immune stimulation from viral infection, autoimmunity, or cancer (Cenerenti et al., 2022). Our work suggests that continuous homeostatic exposure to food and commensals drives a similar CD4^+^ T cell phenotype in the gut. One essential difference between cytotoxic CD4^+^ T cells described in disease states and cytotoxic CD4^+^ T cells induced by steady state intestinal stimulation is expression of CD8αα which has been proposed to raise the threshold for TCR activation (Cheroutre and Lambolez, 2008; Mucida et al., 2013). Therefore, although IE-adapted CD4^+^ CD8αα^+^ T cells are equipped with cytotoxic machinery, they may require stimulation above steady state levels to activate cytotoxic mechanisms. Our data shows that exposure to food protein in tandem with an inflammatory signal (cholera toxin) drives a hyperactive state in tissue-resident IE-adapted CD4^+^ T cells, including upregulation of cytotoxic genes. In Celiac disease, IE-adapted CD4^+^ T cells exhibit highly inflammatory Th1-like responses and contribute to tissue pathology, which is abrogated upon removal of gluten from the diet (Abadie et al., 2012; Costes et al., 2019; Fina et al., 2008). Thus, dysregulation or dysfunction of IE CD4^+^ T cells in addition to Tregs may be one mechanism by which inflammatory responses to harmless gut antigens emerge. However, current functional understanding of cytotoxic CD4^+^ T cells is limited. For example, the extent to which CD4^+^ T cell cytotoxicity depends on cognate antigen presentation via MHC II, as well as the precise cellular targets of CD4-mediated cytolysis, remain unclear (Takeuchi and Saito, 2017). Further work is needed to define whether and how CD4^+^ T cell cytotoxicity contributes to tolerance in the gut or conversely can become dysregulated and contribute to tissue damage.

We show that inflammatory signals disrupt steady state CD4^+^ T cell intestinal adaptation, leading to increased influx of cells skewed towards effector phenotypes. If oral tolerance to a food protein was established prior to inflammatory challenge, then exposure to the same food protein during inflammatory challenge results in clonal expansion of gut Tregs associated with reduced pro-inflammatory CD4^+^ T cell gene expression and protection against food allergy. However, if the primary exposure to a food protein occurs at the same time as the inflammatory stimulus, impaired Treg response and increased pro-inflammatory CD4^+^ T cell gene expression is observed in the gut, associated with development of food allergy. Although pTregs are known to be required for oral tolerance (Garside et al., 1995; Hadis et al., 2011; Mucida et al., 2005), we found that Treg influx during oral tolerance was characterized by increased expression of Helios, a marker typically associated with thymic Tregs which do not respond to intestinal antigen, and reduced *Il10*. Nevertheless, OVA-specific Tregs identified in the tolerance condition were from the *Il10*+ cluster, demonstrating that *Il10*+ pTregs contribute to antigen-specific tolerance in a polyclonal setting whereas the increased Helios+ Tregs may contribute broadly to tissue protection in an antigen-nonspecific manner. While human allergy is typically associated with a Th2 response towards allergens, we and others find the intestinal CD4^+^ T cell response in the OVA/CT mouse model to be skewed towards pro-inflammatory Th17, an effect that has been shown to depend on the microbiota (Zhao et al., 2017). Whether Th17 are effector cells contributing to allergy in C57BL/6 mice or are merely a parallel response to microbiota in the presence of mucosal adjuvant remains unclear.

Our iSell^Tomato^ data demonstrate that important repertoire dynamics accompany T cell differentiation in response to dietary antigens, conclusions previously precluded by the widespread use of TCR monoclonal systems (Mucida et al., 2005; Weiner et al., 2011). Additionally, we provide evidence that steady state exposure to dietary proteins contributes to clonal selection of intestinal CD4^+^ T cells, and identify TCR clones from the tissue that recognize dietary protein. Our findings complement a recent study which used MHC Class II tetramers to show generation of a dietary antigen-specific CD4^+^ T cell response in a natural polyclonal setting, where the primary fate in the LP was Treg differentiation (Hong et al., 2022). Our approach does not rely on epitope-specific discovery, and the clones we identify do not bind the immunodominant OVA epitope previously identified via immunization (Robertson et al., 2000), but rather both bind OVA 26:40. This may suggest distinct dynamics of clonal selection by dietary antigen in the natural oral route. The food antigen-specific clone we identified from steady state feeding derived from IE-adapted or Th1 cells whereas the oral tolerance clone derived from Tregs. Thus, we show that while oral tolerance is characterized by an antigen-specific Treg response, the steady state response can include differentiation to IE-adapted CD4^+^ CD8αα^+^ cells. Food protein can therefore drive diverse phenotypic outcomes in antigen-specific intestinal CD4^+^ T cells including adaptation to the intestinal epithelium.

Altogether, our findings suggest that the prevalent fate of food-responsive CD4^+^ T cells in the steady state intestine is epithelial adaptation, whereas maintenance of oral tolerance in an inflammatory setting correlates with increased influx and clonal expansion of Tregs. We and others have demonstrated an important regulatory activity of epithelium-adapted CD4^+^ T cells in the context of response to diet (Sujino et al., 2016), or in the context of colitis or infection (Basu et al., 2021; Reis et al., 2013). Further, dysregulation of IE CD4^+^ T cells may be one mechanism by which inflammatory responses to harmless gut antigens emerge. Therefore, highly regulated maintenance of epithelium adapted CD4^+^ T cells in addition to Tregs may be critical for preventing inappropriate immune responses to food and subsequent disease.

## Acknowledgements

We thank all Mucida Lab members and Rockefeller University employees for their continuous assistance, particularly: A. Rogoz and S. Gonzalez for the maintenance of mice, RU Genomics core for sequencing, Tri-I Laboratory of Comparative Pathology for histology preparation, K. Gordon, K. Chhosphel, and J.P. Truman for sorting, B. Reis for assistance with figures, and A. Bilate, G. Donaldson, and R. Parsa for critical reading of the manuscript. We also thank the Victora and Lafaille labs for fruitful discussions. This work was supported by The Howard Hughes Medical Institute, R01DK093674, R01DK113375, R21AI144827, and Food Allergy FARE/FASI Consortium.

## Author Contributions

A.L. initiated, designed, performed, and analyzed experiments and wrote the manuscript. A.R., C.H., and M.C.C.C. designed and performed experiments. T.B.R.C performed analysis. D.M. conceived, initiated, designed, and supervised the research and wrote the manuscript. All authors revised and edited the manuscript and figures.

## Competing interests

The authors declare no competing financial interests.

## Materials and Methods

### Animals

Animal care and experimentation were consistent with the NIH guidelines and were approved by the Institutional Animal Care and Use Committee at the Rockefeller University. All mice were maintained at the Rockefeller University animal facilities. Germ-free C57BL/6J mice were obtained from Sarkis Mazmanian and bred and maintained in germ-free isolators. SPF C57BL/6J mice were re-colonized from GF with a single gavage of feces and bred and maintained in SPF conditions. Vertically colonized ex-GF offspring were used for SPF experiments to control for genetic drift in our GF isolators. The Oligo-MM^12^ consortium was a gift from K. McCoy (Univ. Calgary). We colonized GF C57BL/6J breeders with a single gavage of Oligo-MM^12^ and monitored colonization (including the presence of the entire consortium in successive generations) by specific amplification of individual bacterial members by quantitative polymerase chain reaction (qPCR; see below). Oligo-MM^12^ mice were subsequently bred and maintained in isolators and vertically colonized offspring were used for all experiments. *Sell*^Cre-ERT2^ mice were provided by M. Nussenzweig (Merkenschlager et al., 2021), crossed with *Rosa26^CAG-LSL-tdTomato-WPRE^* (007914) mice from Jackson laboratory, and maintained under SPF conditions. CD45.1 OT-II TCR-transgenic mice were originally purchased from Taconic Farms and maintained in our facilities. Mice were used at 8 weeks of age for most experiments, except iSell^Tomato^ where experimental protocols were initiated at 7 weeks of age and endpoint analysis was performed at 11 weeks. Both male and female mice were used for all experiments, except scRNAseq which used exclusively female (for diet) or male (for iSell^Tomato^) mice to avoid sex effects on gene expression.

### Germ-free and Oligo-MM^12^ monitoring

Germ-free status was confirmed by by qPCR analysis using universal 16S rRNA primers (fwd: ACTCCTACGGGAGGCAGCAGT; rev: ATTACCGCGGCTGCTGGC). Colonization of mice by the Oligo-MM^12^ consortium was confirmed and monitored over generations by qPCR, using primer pairs specific to each species as previously described (Nowosad et al., 2020). DNA was extracted from fecal samples using the ZR Fecal DNA kit (Zymo Research) according to the manufacturer’s instructions. Quantitative PCR was performed with the Power SYBR Green master mix (Applied Biosystems). The average cycle threshold (Ct) value of two technical replicates was used to quantify the relative abundance of each species’ 16S ribosomal RNA using the ΔΔCt method, with the universal 16S rRNA primers serving as controls between samples. Relative abundance was corrected according to the genome copy number of 16S rRNA for each species.

### Experimental Diets

For all dietary experiments, breeders were maintained on standard chow diet and breeding cages were switched to amino acid diet when pups were 1 week old to prevent early exposure to food proteins. Mice were subsequently weaned onto experimental diets at 3 weeks and maintained on that diet until endpoint analysis at 8 weeks old unless otherwise indicated. Protein antigen-free solid diet containing free amino acids (Modified TestDiet 9GCV with 5% cellulose; composition details are in Table S1) was irradiated at >45 kGy to ensure sterility for germ-free conditions. GF, Oligo-MM^12^, and ex-GF SPF chow diet control mice were fed autoclaved standard chow diets fortified with extra nutrients to compensate for losses during autoclaving (LabDiet 5K54). For AA+OVA, Ovalbumin grade III (Sigma, A5378) was provided at 1 mg/mL in drinking water, autoclaved for sterility. Casein and Casein-gluten-soy diets were modified from AA diet to contain 50% less amino acid and 50% even mix of casein or casein, gluten, and soy protein (TestDiet 9GU1, 9GU2).

### Serum biochemistry

Serum was collected from 8-week-old mice fed AA or chow diet since weaning after 2 hours of fasting. Serum analysis was performed by IDEXX (USA) using standard protocols.

### Histology

Representative portions of duodenum, ileum, and colon were fixed in 4% paraformaldehyde and embedded in paraffin according to standard protocols. 5-µm sections were mounted on glass slides and stained with hematoxylin and eosin (H&E). Images were acquired on a Keyence BZ-X800 inverted microscope using a 10x 0.3/14.50mm objective lens with brightfield illumination (Keyence). Blinded quantitative evaluation of intestinal pathology was performed according to established methods (Erben et al., 2014). Briefly, each tissue section was microscopically assessed for the extent of inflammatory cell infiltrate (0-7), and changes to the epithelial (0-23) or mucosal (0-15) architecture. The sum of these scores represents the combined pathological score reported for each tissue with 40 being the maximum score and a score <10 indicating normal tissue with minimal to mild inflammation. Further quantification of tissue metrics was performed in ImageJ.

### Isolation of intestinal T cells

Intraepithelial and lamina propria lymphocytes were isolated as previously described (Bilate et al., 2016; Reis et al., 2013). Briefly, small intestines or large intestines (cecum and colon) were harvested and washed in PBS and 1mM dithiothreitol (DTT) followed by 30 mM EDTA. Intraepithelial cells were recovered from the supernatant of DTT and EDTA washes and mononuclear cells were isolated by gradient centrifugation using Percoll. Lamina propria lymphocytes were obtained after collagenase digestion of the tissue.

### Antibodies and flow cytometry analysis

Fluorescent dye–conjugated antibodies were purchased from BD Biosciences, Biolegend, Ebioscience (Thermofisher), or R&D biosciences. The following clones were used: anti-CD4 RM4-5; anti-CD8α 53-6.7; anti-CD8β YTS 156.7.7; anti-CD11b M1/70; anti-CD11c N418; anti-CD44 IM7; anti-CD45 30-F11; anti-CD45.1 A20; anti-CD62L MEL-14, G8.8; anti-CD69 H1.2F3; anti-CD103 2E7; anti-CD117 (cKit) 2B8; anti-F480 BM8; anti-FceR1 MAR-1; anti-Foxp3 FJK-16s; anti-Granzyme B GB11; anti-IL-4 11B11, anti-IL-17A TC11-18H10; anti-Ly6C AL-21; anti-Ly6G RB6-8C5; anti-Nrp1 BAF566; anti-Rorγt Q31-378; anti-Siglec F E50-2440; anti-TCRβ H57-597; anti-TCRγδ GL3; anti-TCR Vα2 B20.1. Live/dead fixable dye Aqua (ThermoFisher Scientific) was used according to manufacturer’s instructions. Intranuclear staining of Foxp3 and intracellular staining of Granzyme B was conducted using Foxp3 Mouse Regulatory T Cell Staining Kit according to kit instructions (eBioscience, USA). For analysis of IL-4 and IL17-A production, cells were incubated at 37C with 100 ng/mL phorbol 12-myristate 13-acetate (PMA, Sigma Aldrich) 200 ng/mL ionomycin (Sigma), and Golgi stop solution containing Monensin (2 mM, BD Biosciences) for 4 hours. Intracellular staining for cytokines was conducted in Perm/Wash buffer after fixation and permeabilization in Fix/ Perm buffer (BD Biosciences, USA) according to kit instructions. Flow cytometry data was acquired on an LSR-II or Symphony flow cytometer (Becton Dickinson, USA) and analyzed using FlowJo software package (Tri-Star, USA). For cell sorting experiments, lymphocytes were sorted on a FACS Aria II instrument as indicated in the figure legends. AccuCheck Counting Beads (Thermo Fisher, USA) were used for counting of absolute cell numbers. For flow cytometric analysis the following gating strategies were used to identify cell populations. T cells: single, live, lymphocytes (based on FSC, SSC and live/dead fixable dye Aqua stain), CD45+, TCRgδ+ (for gδ T cells) or TCRβ+ (for all other T cells); CD4^+^ T cells: CD4+, CD8β-. CD8αβ^+^ T cells: CD4-, CD8β+. CD8αα^+^ T cells: CD4-, CD8β-, CD8α+. Myeloid cells: single, live, CD45+; Eosinophils: Siglec F+, CD11b+; Mast cells: cKit+, FCer1+; Macrophages: CD11b+, CD11c+, F480+; Monocytes: CD11b+ CD11c-Ly6c+; Neutrophils: Ly6G+ CD11b+.

### OTII Transfer Experiment

Naïve CD4 T cells from spleen and lymph nodes of CD45.1 OTII TCR transgenic mice were isolated by negative selection using biotinylated antibodies against CD8α, CD25, CD11c, CD11b, TER-119, NK1.1, and B220 and anti-biotin MACS beads (Miltenyi Biotec). 1x10^6^ cells were transferred by retro-orbital injection to CD45.2 hosts under isoflurane gas anesthesia. Host mice were provided regular drinking water, or drinking water supplemented with 1 mg/mL Ovalbumin grade III (Sigma, A5378). After 48 hours, mesenteric lymph nodes were collected and CD45.1+ TCRVα2+ CD4+ OTII cells were analyzed for activation by expression of CD69.

### Tamoxifen Treatment

Tamoxifen (Sigma) was dissolved in corn oil (Sigma) and 10% ethanol, shaking at 37°C for 30 min-1 h. Two doses of Tamoxifen (5 mg/dose) were administered to mice via oral gavage at 50 mg/mL, 3 days and 1 day before start of treatment protocol.

### OVA/Cholera Toxin Allergy Model

Ovalbumin (OVA) grade III (Sigma, A5378) was provided at 0.1% in drinking water, autoclaved for sterility, for 3 days (Feeding, Tolerance) or not (Allergy) to initialize tolerance. All mice were then provided regular drinking water for 1 week. 1 mg OVA in 0.2M sodium bicarbonate (Feeding), or 1 mg OVA + 20 ug cholera toxin (List Biological 100B) in 0.2M sodium bicarbonate (Tolerance, Allergy), were provided once per week for 3 weeks, followed by endpoint analysis 2 days after the final dose. Serum was harvested for ELISA one day prior to endpoint analysis. For anaphylaxis experiments, the above protocol was followed except for the following modifications: OVA or OVA/CT was provided once per week for 4 weeks, followed by challenge 7 days after the final dose. Implantable electronic temperature probes (Avidity IPTT-300) were injected s.c. 1 day prior to challenge. Mice were challenged with 5 mg OVA i.p. and body temperature was measured every 10 minutes for 50 minutes. Mice used in anaphylaxis experiments were not used for downstream sequencing analysis.

### ELISA

Lipocalin-2 was analyzed by using Lcn-2 ELISA kit (R&D, MN) as described by (Chassaing et al., 2012). IgE and OVA specific IgG1 ELISAs were performed as described previously (Atarashi et al., 2011).

### 16S rRNA sequencing

Intestinal contents were collected fresh from the whole small intestine or cecum at endpoint analysis. Littermate controls were used for all 16S experiments to control for maternal effects. Samples were collected from at least four different cages per experimental group to control for cage effect. Samples were prepared for 16S rRNA sequencing following the 16S Illumina Amplicon protocol from the Earth Microbiome Project (Caporaso et al., 2011). Libraries were sequenced using Miseq 2x150 using a 15% PhiX spike. Sequence processing and analysis was performed in R. Briefly, read assembly into amplicon sequence variants (ASVs) and taxonomic assignment on the Silva database were performed using Dada2 (v.1.2.6). Taxa not seen more than 10 times in at least 20% of samples were removed from analysis. ASV quantification and analyses Tawere performed using Phyloseq (v1.42) (McMurdie and Holmes, 2013).

### Single cell RNA-seq library preparation

Lymphocytes isolated from the small intestine epithelium or lamina propria were isolated as described above and indexed with TotalSeqC Hashtag (BioLegend) cell surface antibodies, with 2 barcodes used per sample for deeper multiplexing. Total CD4^+^ T cells were sorted, pooled, and immediately loaded onto a Chromium Controller (10x Genomics). For Sell^Tomato^ sorted cells were pooled at a ratio of 1:3 Tomato^+^:Tomato^−^. 5’ Gene expression, VDJ, and Feature Barcode libraries were prepared using the Chromium Single Cell 5′ v2 Reagent Kit (10x Genomics) according to manufacturer’s protocol at the Genomics core of Rockefeller University. Libraries were sequenced on a NovaSeq. Hashtag indexing was used to demultiplex the sequencing data and generate gene-barcode matrices.

### Data processing of single cell RNAseq and single cell TCRseq libraries

Raw fastq files from our 10x libraries were processed with Cellranger count (v6.2.0) using the 10x Genomics prebuilt mouse reference (v3.0.0 mm10) (for diets) or a customized mouse genome (mm10) that included the ai9Tomato sequence plasmid as an artificial chromosome with the Ai9Tomato and STOP sequences annotated as features (for Sell^Tomato^). Analyses were performed in R 4.2.2 (R Core Team, 2018). Quality control was performed by removing cells with high (> 10% for the antigen-free diet libraries, >5% for Sell^Tomato^) mitochondrial unique molecular identifier (UMI) content. Cells not expressing *Trac* or *Cd4* were excluded from our analysis. We defined Tomato^+^ and Tomato^−^ cells post-sequencing using normalized UMI counts for *tdTomato* and *Stop* (Tomato^+^: *tdTomato* > *Stop*. Tomato^−^: *tdTomato* < *Stop*), discarding ambiguous cells where *tdTomato* = *Stop*. The matrix of UMI counts was normalized by applying a regression model with a negative binomial distribution, available through the SCTransform function in the Seurat (v41.-4.3.) package (Hafemeister and Satija, 2019). The top 3000 variable genes were first used for dimensional reduction by PCA using the scaled data. The first 30 principal components were further used for visualization using the Manifold Approximation and Projection (UMAP) and cell clustering (Hafemeister and Satija, 2019; Stuart et al., 2019). TCR contigs and annotation were performed with the Cellranger vdj workflow from 10x Genomics and the prebuild mouse reference (v3.1.0 mm10). Contigs filtering, clonotype calling, and downstream TCR analysis was performed using scRepertoire (v1.5.2) (Borcherding et al., 2020). Further processing, statistical analysis, and visualizations were performed using ggplot2 (v3.4.1) (Wickham, 2016) and rstatix (v0.7.2).

### Signature Scoring

IE signature scores were calculated using UCell (v2.2) (Andreatta and Carmona, 2021) using the following genes as input: *Nkg7, Ccl5, Cd160, Itgae, Gzma, Gzmb, Cd7, Prf1, Lag3, Cd8a*.

### 3D Volcano

3D volcano plots were generated using volcano3D (v1.2-2.0.8) (Lewis et al., 2019). Briefly, genes differentially expressed across 3 groups were identified by FDR-adjusted Kruskal-Wallis test (p-adj <0.05) considering only genes expressed in at least 10% of cells in any group. Upregulation in each group was further determined by FDR-adjusted Wilcoxon rank sum test (p-adj <0.05, Log2FC >0.5). Each point on the plot represents a gene colored by the group(s) in which they are significantly upregulated, or uncolored for nonsignificant genes. Distance from the origin for each gene represents a z-score calculated based on -log10 p-values from Kruskal-Wallis comparisons between all 3 groups, with genes further from the origin having higher significance. Degree of upregulation of a gene in a given condition is indicated by its angle on the plot relative to condition-labeled axes. Scaled gene expression data was used for all calculations except fold change. Select genes of interest are labeled on each plot.

### Circos Plots

TCR sharing (clonal overlap) was visualized using Circos to create circular plots aesthetics (Krzywinski et al., 2009). Each segment denotes a mouse as indicated in figures, with bands representing clonal sharing based on paired TCRα and TCRβ amino acid sequences. NKT cells and cells without paired TCRα and TCRβ sequenced were not considered in analysis.

### Statistical Analyses

Statistical analysis was carried out using GraphPad Prism v.9. Flow cytometry analysis was carried out using FlowJo software. Data in graphs show mean +/-SEM and p values <0.05 were considered significant. Repertoire diversity was analyzed by Diversity 50 (D50) calculated in R as the fraction of dominant clones that account for the cumulative 50% of the total paired CDR3s. GraphPadPrism v.9 was used for graphs and Adobe Illustrator 2021 used to assemble and edit figures.

### TCR Hybridoma Generation

Select TCRs from the scRNAseq datasets were synthesized (TCRα and TCRβ linked by P2A) and cloned into pMSCV-mCD4 vectors (Twist Bioscience). Phoenix-Eco cells (Addgene CRL-3214) were used for retrovirus production. Phoenix were grown to 60-80% confluence in DMEM in 10 cm culture dishes pre-coated 2 mL 0.01% poly-L-lysine (Sigma A-005-M) in 8 mL 0.1% gelatine (Sigma 1040700500) and then transfected with a mix containing 5 ug pMSCV-mCD4-TCR plasmid, 2 ug pEco (Addgene 12371), 72 uL 1 mg/mL polyethylenamine (Polysciences 23966), and 600 uL DMEM. Media was changed at 24h and harvested at 48h for viral transduction. NFAT-GFP hybridoma cells were plated 1x10^6^/mL in viral supernatant with 5 ug/mL polybrene (Sigma TR-1003-G) in 6-well plates pre-coated with 15 ug retronectin (Takara T100A). Plates were spinfected by centrifugation at 2500 rpm 32C for 90 minutes. Cells were then cultured for 2-7 days in IMDM + 10% FBS + Pen/Strep/L-glutamine + 50 uM β-me + 1 mM Sodium Pyruvate before FACS selection for TCR+ CD3+ cells.

### TCR testing for OVA specificity

Dendritic cells were collected from mouse spleens (Miltenyi 130-092-465) and co-cultured 5x10^5^/mL with 50 ug OVA grade III (Sigma A5378) for 4h at 37C in 96-well U-bottom plates to load with antigen, or left unloaded for negative control. 2.5x10^4^ TCR-expressing NFAT-GFP cells were added to each well and incubated overnight. For positive control, 0.2 uL anti-CD3 (BD 553057) was added to a well with NFAT and unloaded DCs. NFAT-GFP response to TCR stimulation was measured by flow cytometry. To test epitope specificity, crude 15 aa peptides covering the length of OVA with a 5aa shift were synthesized (LifeTein) and used in the assay as above at 1 uM per well. TCR stimulation was measured by IL-2 ELISA on culture supernatant (BD 555148).

## Supplemental Information

Supplemental Figure Legends S1-S5

Figures S1-S5

Tables S1-S12

## Supplemental Figure Legends

**Figure S1 AA diet mice are heathy and display no evidence of intestinal damage or inflammation. (A)** Percent of original body weight (top) or fecal lipocalin-2 levels measured by ELISA (bottom) from SPF mice at indicated weeks after weaning onto AA or standard chow diet. Mean ± SEM representative of 2 independent experiments using 19-22 mice per group. Comparisons between diets within timepoints are not significant as calculated by unpaired t test. **(B)** Serum nutritional biomarkers measured in 8-week-old SPF mice fed AA or chow diet since weaning. ALP – alkaline phosphatase, AST – aspartate aminotransferase, BUN – blood urea nitrogen, HDL – high density lipoprotein, LDL – low density lipoprotein, LDH – lactose dehydrogenase. 2 independent experiments, each point represents pooled serum from 4-6 mice. **(C-F)** SPF or GF mice were fed AA or standard chow diet from weaning until analysis at 8 weeks old. **(C)** Small intestine length. **(D)** Representative H&E histology images with 200 μm scale bar. **(E)** H&E pathology scores based on 1 image per tissue per mouse where 40 is the maximum score (top left) or tissue morphology measures where each dot represents the average of 4 measurements per tissue per mouse. **(F)** Flow cytometry analysis of myeloid cells in the small intestine IE and LP. Mean ± SD representative of 2 independent experiments with 6-9 mice per group. Unpaired t tests with Holm-Sidak multiple comparison test, p-values <0.05 labeled on plot.

**Figure S2 Dietary signals promote accumulation and adaptation of intestinal CD4^+^ T cells in the small intestine epithelium – additional data supporting Figures 1 & 2. (A-D)** Flow cytometry from the small intestine (A-C) or large intestine (D) IE or LP of 8-week-old SPF mice weaned onto AA or standard chow diet measuring frequency or absolute count of the indicated cell subsets. Unpaired t-tests with p-values <0.05 shown on each plot. Mean ± SD representative of 3-5 independent experiments using 7-18 mice per group. **(E)** Flow cytometry of transferred OTII CD4^+^ T cells from the mesenteric lymph nodes (mLN) after 48 hours of OVA supplied 1 mg/mL in drinking water as indicated. One-way ANOVA with Tukey multiple comparison test, p-values <0.05 labeled on plot. Data is representative of 2 independent experiments with 3-4 mice per group. **(F-H)** Flow cytometry from the small intestine IE or LP of 8-week-old SPF mice weaned onto AA diet with or without 1 mg/mL OVA supplied in drinking water (F-G) or AA, casein, or casein-gluten-soy diet (H) measuring frequency or absolute count of the indicated cell subsets. Unpaired t-tests (F-G) or one-way ANOVA with Tukey multiple comparison test (H) with p-values <0.05 shown on each plot. Mean + SD representative of 2-3 independent experiments using 3-11 mice per group. **(I-J)** 16S rRNA sequencing of cecum contents of 8-week-old SPF mice fed AA or standard chow diet represented by relative phyla abundance (I), and SI Chao1 alpha diversity with mean ± SD and unpaired t-test (J). Data is from 4 independent experiments using 11-15 mice per condition. **(K-L)** Flow cytometry from the IE or LP of 8-week-old GF or Oligo-MM^12^ mice weaned onto AA or standard chow diet measuring frequency or absolute count of the indicated cell subsets. Dashed lines show mean value from SPF Chow (red) or SPF AA (blue). Two-way ANOVA with p-values beneath each plot and Holm-Šidák multiple comparison test between diets within each colonization represented by p-values within each plot. Only p-values <0.05 from the multiple comparison test are shown. Barplots show mean + SD representative of 2-3 independent experiments using 6-12 mice per group.

**Figure S3 Chow diet promotes microbiota-independent epithelial adaptation and cytotoxic transcriptional programming of intestinal CD4+ T cells – additional data supporting Figures 2 & 3. (A-E, G-J)** CD4+ T cells were sorted from the IE or LP of 8-week-old GF or Oligo-MM12 mice weaned onto AA, AA+OVA, or standard chow diet and single cell RNA sequencing was performed using the 10X Genomics platform, pooling 2-4 mice per diet/colonization group. The data shown is for all sequenced CD4+ T cells (A-E) or subclustered Tregs (G-J) **(A)** Number of cells sequenced per indicated sample, colored by sequencing batch (left), and violin plots showing number of detected RNA molecules, number of sequenced genes, or percent mitochondrial DNA per cell per sequencing batch (right). **(B, G)** Top 5 differentially expressed genes (ranked by fold change) in each UMAP gene expression cluster from total CD4+ T cells (B) or Tregs (G). Wilcoxon rank sum test (p<0.01). **(C, J)** Frequency of mature clusters (IE1, IE2, IE3, and Th1 combined; C) or Il10 high Tregs (J). Two-way ANOVA with p-values beneath each plot. **(D, I)** IE signature score of total CD4+ T cell (E) or Treg (I) gene expression clusters. **(E)** Frequency of IE2 or IE3 out of total IE CD4+ T cells. one-way ANOVA with Tukey multiple comparison test p-values <0.05 displayed on plot. **(F)** Flow cytometry from the IE or LP of 8-week-old GF or Oligo-MM12 mice weaned onto AA or standard chow diet measuring frequency or absolute count of the indicated cell subsets. Dashed lines show mean value from SPF Chow (red) or SPF AA (blue). Two-way ANOVA with p-values beneath each plot and Holm-Šidák multiple comparison test between diets within each colonization represented by p-values within each plot. Only p-values <0.05 from the multiple comparison test are shown. Barplots show mean + SD representative of 2-3 independent experiments using 3-9 mice per group. **(H)** Frequency of IE subclusters in IE or LP. **(K)** Flow cytometry from the IE of 8-week-old SPF mice weaned onto standard chow diet measuring frequency of Granzyme B out of the indicated cell subsets. one-way ANOVA with Tukey multiple comparison test p-values <0.05 displayed on plot. Mean + SEM representative of 3 independent experiments using 8 mice per group.

**Figure S4 Intestinal CD4^+^ T cell responses during OVA feeding, tolerance, or allergy – additional data supporting Figures 6 and 7**. iSell^Tomato^ mice were analyzed on day 26 after treatment with tamoxifen to permanently label naïve T cells and then exposure to OVA in the context of feeding, tolerance, or allergy. **(A)** Representative H&E histology images with 200 μm scale bar (left) and pathology scores based on 1 image per tissue per mouse where 40 is the maximum score (right). Mean + SD representative of 2 independent experiments using 4-5 mice per group. **(B-D)** Flow cytometry measuring frequency of Tomato^+^ CD4^+^ T cells in the large intestine (B) or of the indicated CD4^+^ T cell subsets out of Tomato^+^ or Tomato^-^ CD4^+^ T cells in the IE (C) or LP (D). One-way ANOVA with Tukey multiple comparison test, p-values <0.05 labeled on plot. Data is representative of 2 independent experiments with 5-8 mice per group. **(E-I)** scRNAseq of 11,217 Tomato^+^ and Tomato^-^ CD4^+^ T cells from the IE and LP using 4-5 mice per condition pooled across 2 independent experiments and sequencing runs. (**E**) Captured cells per sample in 10X sequencing experiment with Tomato^+^ or Tomato^−^ assignments. **(F)** Violin plots showing number of detected RNA molecules, number of sequenced genes, or percent mitochondrial DNA per cell per sequencing run. (**G**) Top 5 differentially expressed genes (ranked by fold change) in each UMAP gene expression cluster from total CD4^+^ T cells. Wilcoxon rank sum test (p<0.01). (**H**) UMAP visualization of sequenced cells positioned by gene expression similarity and colored by tissue (top left) or treatment group (top right) or Tomato assignment (bottom left).

**Figure S5 Intestinal CD4+ T cell responses and antigen specificity during OVA feeding, tolerance, or allergy – additional data supporting Figures 7 and 8. (A-G)** scRNAseq of 11,217 Tomato+ and Tomato-CD4+ T cells from the IE and LP of mice on day 26 of OVA feeding, tolerance, or allergy protocols using 4-5 mice per condition pooled across 2 independent experiments and sequencing runs. **(A-B)** Expression (Pearson residuals) of hallmark Th2 genes (A) or IE or IE4 signature genes (B) in the indicated cell clusters. **(C)** Frequency of Tomato labeling within each gene expression cluster. **(D, F)** Frequency of the indicated cell subsets within each group displaying p<0.05 from one-way ANOVA with Tukey multiple comparison test. **(E)** 3-way volcano plot showing differential gene expression between conditions in Tomato-CD4+ T cells from the IE. Colored genes are differentially expressed (p-adj <0.05 from FDR-corrected Kruskal-Wallis Test and log2 fold change >0.5), colored by the condition(s) in which they are upregulated. Select genes of interest are labeled on each plot. **(G)** Top 5 differentially expressed genes (ranked by fold change) in each Treg subcluster. Wilcoxon rank sum test (p<0.01). **(H)** Overlapping OVA peptide library to determine epitope specificity of OVA-responsive TCRs. A library of 15 amino acid (aa) OVA peptides with 10 aa overlap and 5 aa shift covering the full length of OVA were tested for TCR response in NFAT hybridomas expressing candidate TCRs. Response was measured with an IL-2 ELISA and data is represented as fold increase in IL-2 production compared to the positive control (a-CD3).

## Notes

### Competing Interest Statement

The authors have declared no competing interest.

