## Supplementary figures and images for "Dietary protein shapes the profile and repertoire of intestinal CD4^+^ T cells"

### Figure S1

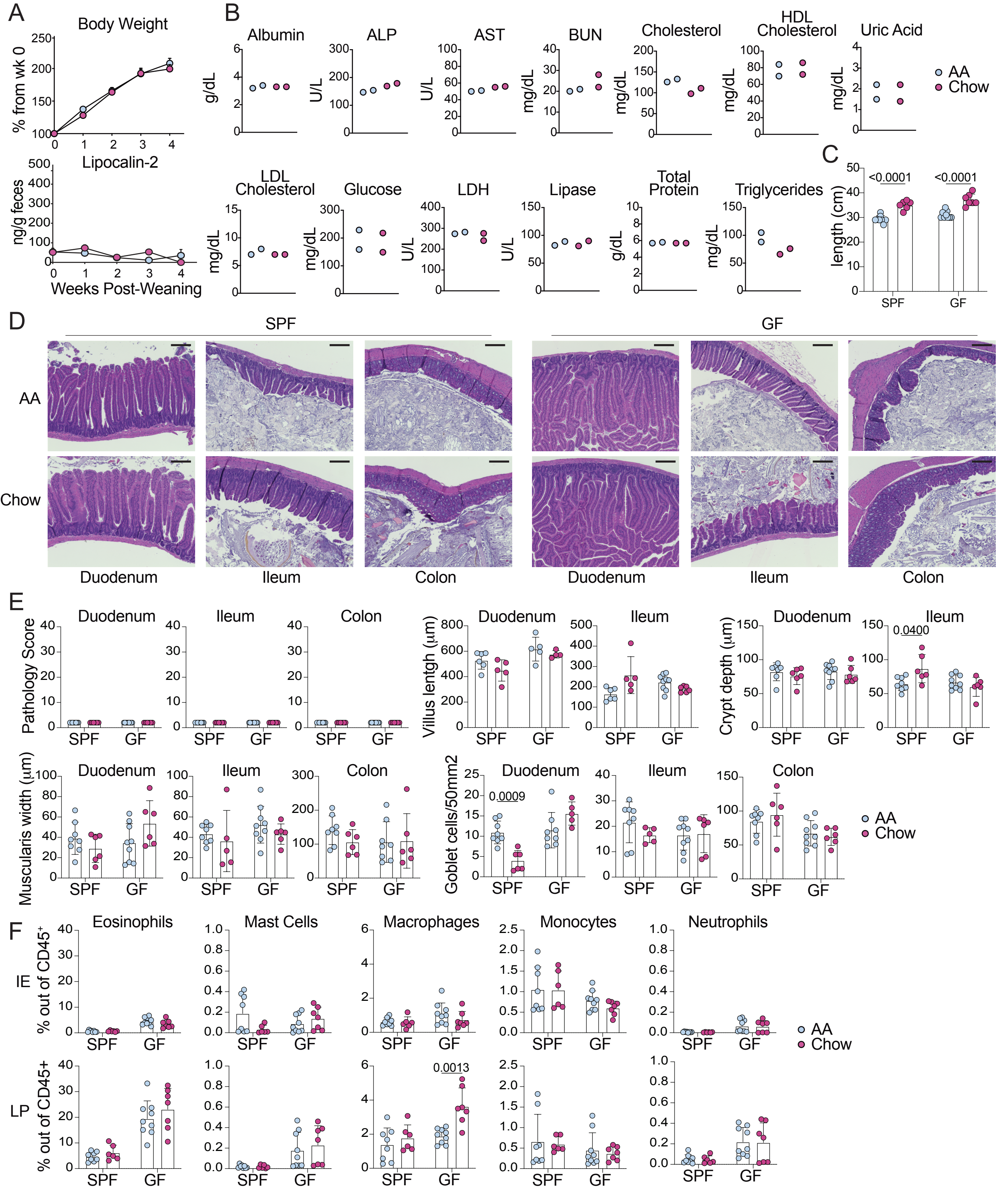

### Figure S2

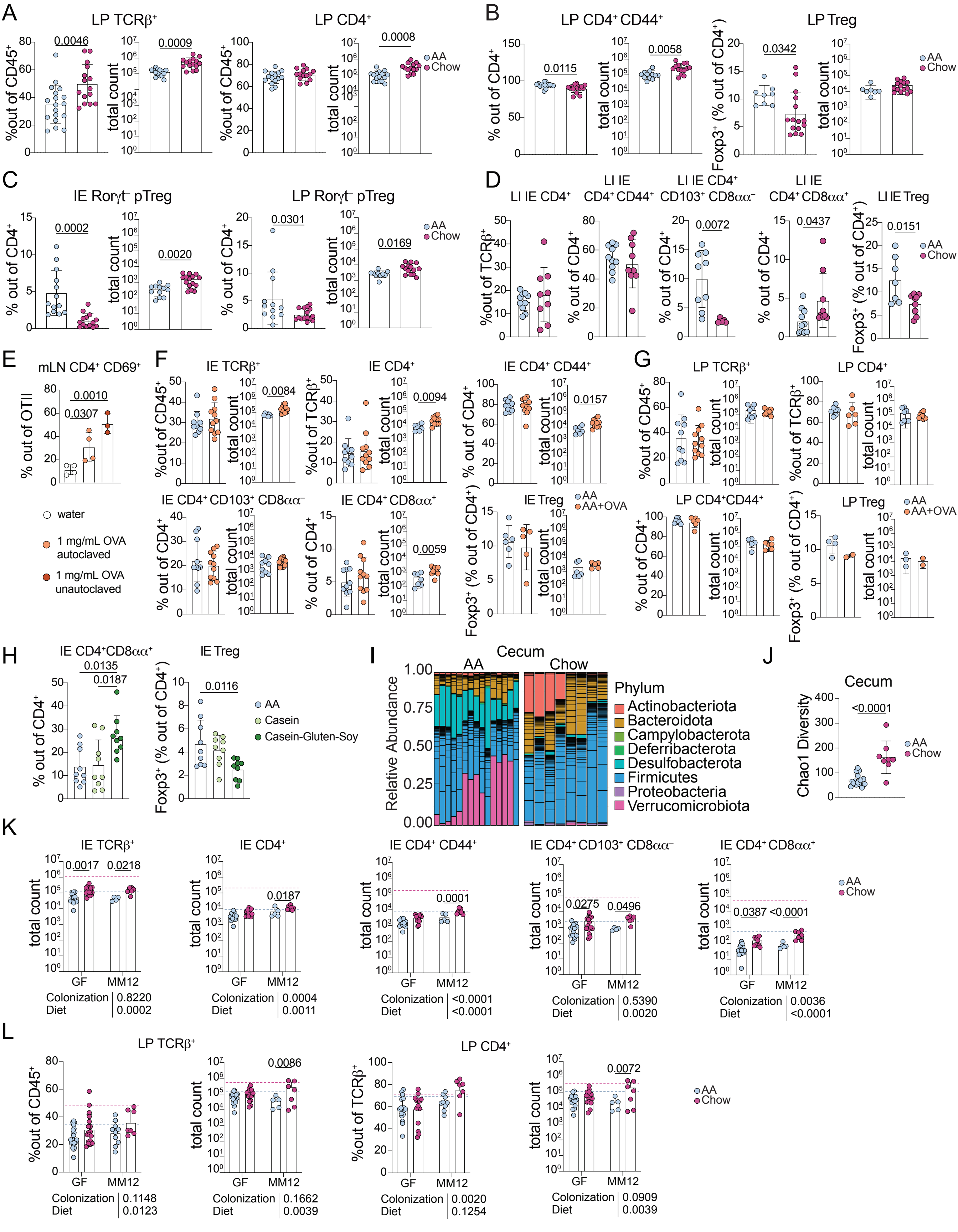

### Figure S3

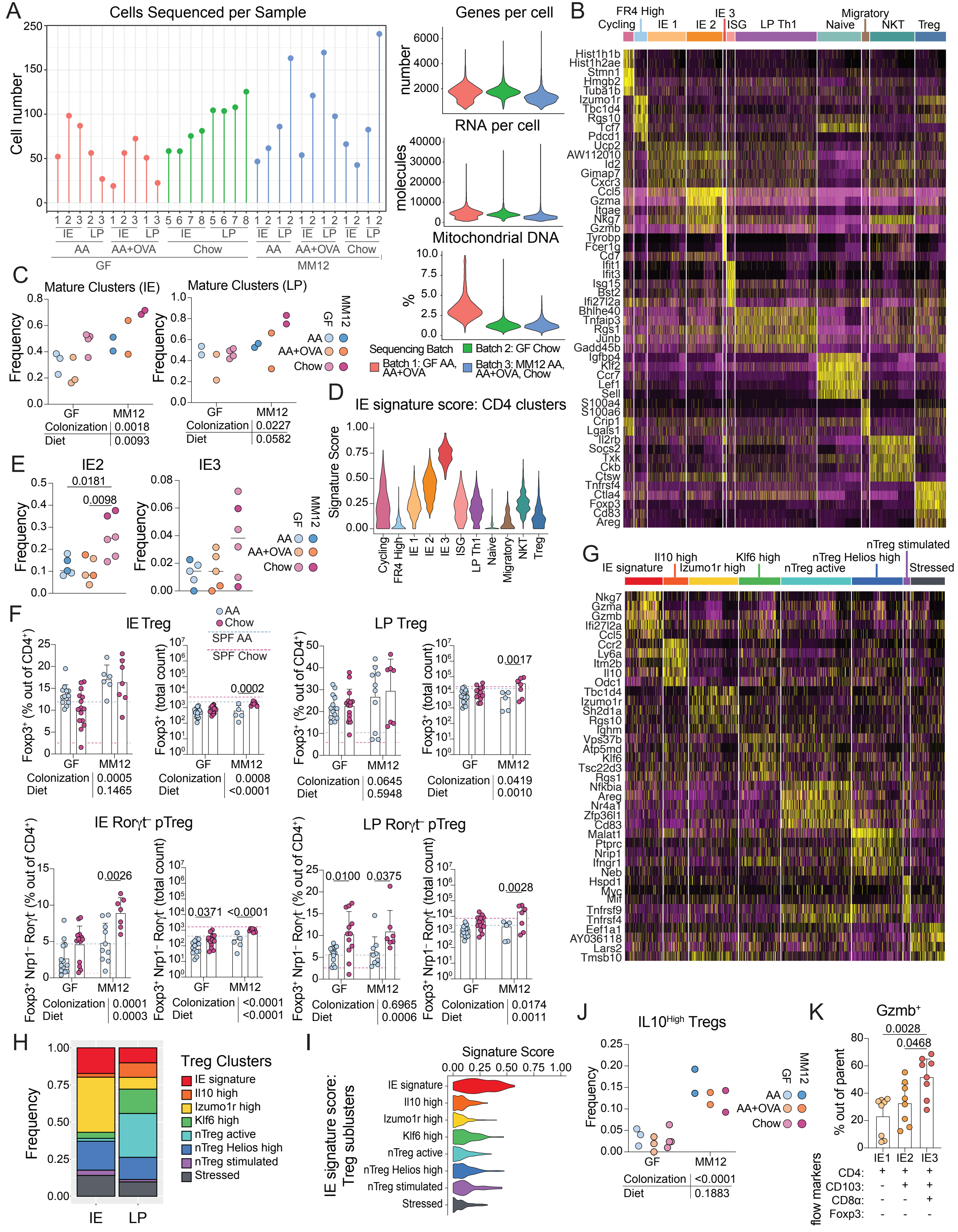

### Figure S4

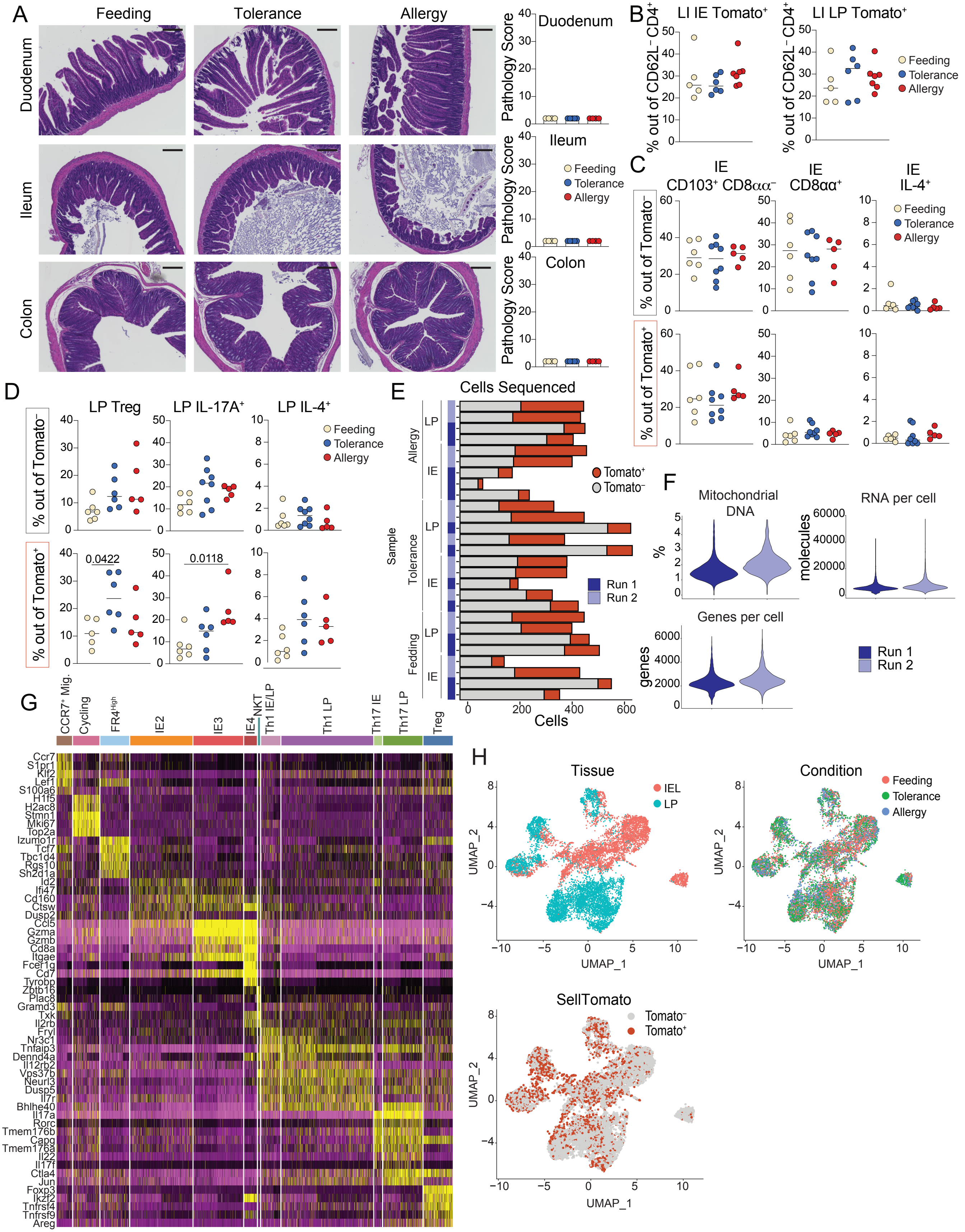

### Figure S5

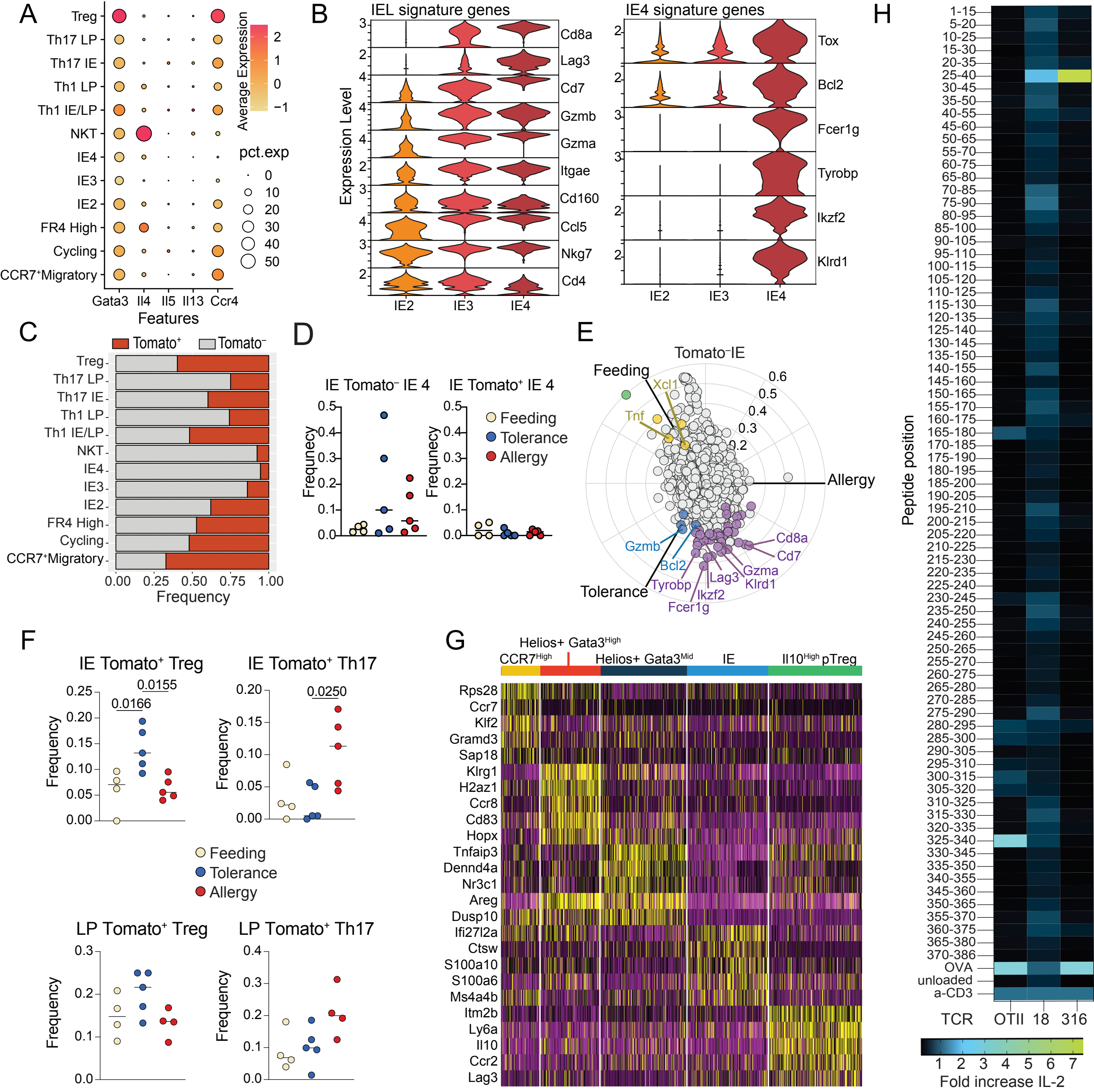
