## Supplementary material for "Dietary protein shapes the profile and repertoire of intestinal CD4^+^ T cells": Table S1

**Table S1.** Composition of protein antigen-free solid diet (AA).

**Ingredients (%)**

Corn starch (35.71), Sucrose (25.90), Amino Acid Premix (17.57), Mineral Premix (10.00), Corn oil (5.00), Powdered cellulose (5.00), Sodium bicarbonate (1.00), Vitamin premix (0.20), Choline chloride (0.10), Ethoxyquin (0.02), DL-Alpha Tocopheryl Acetate (0.004)

| Category | Component | % in Diet |
|---|---|---|
| **Protein** | | **15.00** |
| | Arginine | 0.58 |
| | Histidine | 0.42 |
| | Isoleucine | 0.79 |
| | Leucine | 1.43 |
| | Lysine | 1.20 |
| | Methionine | 0.42 |
| | Cystine | 0.06 |
| | Phenylalanine | 0.79 |
| | Tryosine | 0.83 |
| | Threonine | 0.64 |
| | Tryptophan | 0.18 |
| | Valine | 0.94 |
| | Alanine | 0.45 |
| | Aspartic Acid | 1.06 |
| | Glutamic Acid | 3.37 |
| | Glycine | 0.32 |
| | Proline | 1.95 |
| | Serine | 0.91 |
| | Taurine | 0.00 |
| **Fat** | | **5.10** |
| | Cholesterol | 0 |
| | Linoleic Acid | 2.86 |
| | Linolenic Acid | 0.05 |
| | Arachidonic Acid | 0.00 |
| | Omega-3 Fatty Acids | 0.05 |
| | Total Saturated Fatty Acids | 0.64 |
| | Total Monounsaturated Fatty Acids | 1.21 |
| | Polyunsaturated Fatty Acids | 2.90 |

| | | |
|---|---|---|
| **Fiber** | Cellulose | **5.00** |
| **Carbohydrates** | | **66.20** |
| **Minerals** | | |
| | Calcium | 1.21 |
| | Phosphorous | 0.72 |
| | Potassium | 0.41 |
| | Magnesium | 0.01 |
| | Sodium | 0.64 |
| | Chloride | 1.1 |
| | Fluorine | 0.0 ppm |
| | Iron | 87 ppm |
| | Zinc | 52 ppm |
| | Manganese | 211 ppm |
| | Copper | 5.0 ppm |
| | Cobalt | 0.3 ppm |
| | Iodine | 30.58 ppm |
| | Chromium | 0.0 ppm |
| | Molybdenum | 35.69 ppm |
| | Selenium | 0.46 ppm |
| **Vitamins** | | |
| | Vitamin A | 5.2 IU/g |
| | Vitamin D-3 | 0.9 IU/g |
| | Vitamin E | 20.0 IU/kg |
| | Vitamin K | 2.00 ppm |
| | Thiamin | 18.4 ppm |
| | Riboflavin | 10.0 ppm |
| | Niacin | 50 ppm |
| | Pantothenic Acid | 28 ppm |
| | Folic Acid | 4.0 ppm |
| | Pyridoxine | 4.9 ppm |
| | Biotin | 0.6 ppm |
| | Vitamin B12 | 38 mcg/kg |
| | Choline Chloride | 700 ppm |
| | Ascorbic Acid | 250.0 ppm |
| **Energy** | | 3.7 kcal/g |
